# Field microenvironments regulate crop diel transcript and metabolite rhythms

**DOI:** 10.1101/2021.04.08.439063

**Authors:** Luíza Lane Barros Dantas, Maíra Marins Dourado, Natalia Oliveira de Lima, Natale Cavaçana, Milton Yutaka Nishiyama, Glaucia Mendes Souza, Monalisa Sampaio Carneiro, Camila Caldana, Carlos Takeshi Hotta

## Abstract

- Most research in plant chronobiology was done in laboratory conditions. However, they usually fail to mimic natural conditions and their slight fluctuations, highlighting or obfuscating rhythmicity. High-density crops, such as sugarcane (*Saccharum* hybrid), generate field microenvironments with specific light and temperature due to mutual shading.
- We measured the metabolic and transcriptional rhythms in the leaves of 4-month-old (4 mo) and 9 mo sugarcane grown in the field. Most of the assayed rhythms in 9 mo sugarcane peaked >1 h later than in 4 mo sugarcane, including rhythms of the circadian clock gene, *LATE ELONGATED HYPOCOTYL* (*LHY*).
- We hypothesized that older sugarcane perceives dawn later than younger sugarcane due to self-shading. As a test, we measured *LHY* rhythms in plants on the east and the west side of a field. We also tested if a wooden wall built between lines of sugarcane changed their rhythms. The *LHY* peak was delayed in the plants in the west of the field or beyond the wall; both shaded at dawn.
- We conclude that plants in the same field may have different phases due to field microenvironments, impacting important agronomical traits, such as flowering time, stalk weight and number.

## Introduction

Plants are sessile organisms living in constantly changing environments. Some of those changes are rhythmic due to the movements of the tilted Earth around the sun, bringing seasons, and around itself, bringing day and night. The circadian oscillator is an adaptation for life in rhythmic environments. It is an internal regulatory network that allows plants to track the time of the day by generating responses on both metabolism and physiology levels. The ability to anticipate the rhythmic changes in the environment increases plants fitness (Green *et al*., 2002; Dodd *et al*., 2005). Plants that cannot keep their rhythms desynchronize with the environment and assimilate less carbon (C), accumulate less biomass and have lower water use efficiency (Dodd *et al*., 2005).

The circadian oscillator network comprises a central oscillator, input pathways and output pathways. The central oscillator generates rhythms independently of environmental cues (zeitgebers), such as light and temperature. Input pathways continuously feed the central oscillator with internal and external information, synchronizing it with environmental rhythms (Webb *et al*., 2019). Output pathways gather temporal information from the interactions between the central oscillator and the input pathways and translate it into timely regulated metabolic and physiologic responses. The plant central oscillator includes several interlocked feedback loops based on the regulation of transcription and translation. For example, dawn is marked by an increase in transcripts of *CIRCADIAN CLOCK ASSOCIATED 1* (*CCA1*, not found in monocots), *LATE ELONGATED HYPOCOTYL* (*LHY*) and *REVEILLE 8* (*RVE8*) expression (Alabadí *et al*., 2001; Rawat *et al*., 2011; Gray *et al*., 2017), which in turn regulate and are regulated by the expression of members of the PSEUDO RESPONSE REGULATOR (PRR) gene family during the day (Nakamichi *et al*., 2010; Huang *et al*., 2012). In monocots, this includes *PRR1*, *PRR37*, *PRR59*, *PRR73*, and *PRR95* (Hotta *et al*., 2013; Calixto *et al*., 2015; Dantas *et al*., 2020). As LHY expression declines during the day, the expression of PRR1, usually called TIMING OF CAB EXPRESSION (TOC1), increases, leading to a peak near dusk (Alabadí *et al*., 2001). In *Arabidopsis thaliana* (L.) Heynh., TOC1 degradation is regulated by an interplay between ZEITLUPE (AtZTL) and GIGANTEA (AtGI) (Kim *et al*., 2007; Cha *et al*., 2017). In eudicots, a protein complex called EVENING COMPLEX (EC) is composed of LUX ARRHTHMO (LUX), EARLY FLOWERING 3 and 4 (ELF3 and ELF4) and is assembled during the night, repressing many other central oscillator genes (Herrero *et al*., 2012). The EC still needs to be confirmed in monocots, even though *ELF3* is present and functional (Zhao *et al*., 2012; Huang *et al*., 2017).

Experiments done in laboratory conditions frequently fail to fully mimic natural conditions, especially their rhythms (Annunziata *et al*., 2017; Song *et al*., 2018). For example, in rice (*Oryza sativa* L.), mutations in homolog *OsGI* lead to a delay in flowering in short days and long days in laboratory conditions. However, this phenotype was not observed in long days in the field (Izawa *et al*., 2011). In *Arabidopsis*, flowering under artificial conditions does not fully mimic flowering in nature due to the light quality and spectra and the temperature gradients found in natural conditions (Song *et al*., 2018). *Arabidopsis* that was grown under artificial light also had different metabolic profiles than plants grown under natural light (Annunziata *et al*., 2017). Rhythms in the transcription of plants grown in natural conditions or the field are regulated by the circadian oscillator and by changes in external conditions, such as temperature, light intensity, humidity, and photoperiod, or internal conditions, such as plant age and plant physiology (Nagano *et al*., 2012, 2019; Matsuzaki *et al*., 2015; Panter *et al*., 2019; Dantas *et al*., 2020). A better understanding of plant rhythms in the field is essential to translate this knowledge to agriculture (Steed *et al*., 2021).

Here we have measured metabolic and transcriptional rhythms in sugarcane leaves grown in the field. We show that transcript and metabolite rhythms in 4-months-old (4 mo) sugarcane have earlier peaks in the day than 9 mo sugarcane, with phase shifts ranging from 2 h to 12 h. These phase shifts are correlated with changes in the peak of expression of circadian oscillator genes. Such variation in the timing of the peaks was confirmed by experiments done on plants grown on the east and west sides of the field or on plants that were separated by a wooden wall. We conclude that external rhythms in field microenvironments regulate metabolite rhythms and transcripts associated with the sugarcane circadian oscillator.

## Materials and Methods

### Field conditions, plant growth, and harvesting

All three assayed sugarcane fields were located in the same area, at the Federal University of São Carlos, campus Araras, in São Paulo state, Brazil (22°21′25″ S, 47°23′3″ W; altitude of 611 m). The soil was classified as a Typic Eutroferric Red Latosol. Sugarcane from the commercial variety SP80 3280 (*Saccharum* hybrid) was used in all experiments. The environmental conditions for the three experiments were collected from a local weather station or from within the field using a handheld lux meter and a thermometer at the height of the leaves (Fig. S1). Light intensity was measured with the sensor parallel to the ground. Plants were sampled every 2 h for 26 h, starting 1.5 h before dawn. The leaves +1 of nine sugarcane individuals were harvested and separated into three pools for all experiments, each with three individuals, and each pool was considered a biological replicate. The leaves +1 are the first fully photosynthetically active leaf in sugarcane. Each sampling took less than 30 min, and the harvested tissue was immediately frozen in liquid nitrogen. In the first experiment, sugarcane was planted in April/2012 and sampled in August/2012 (4-months-old), during winter, and in January/2013 (9-months-old), during summer. In the winter harvesting, dawn was at 6:30, and dusk was 18:00 (11.5 h day/12.5 h night). In summer harvesting, dawn was at 5:45, and dusk was 19:00 (13.3 h day/10.7 h night). For the second experiment (east and west margins), plants were planted in October/2014 and harvested in March/2015 (5 months old) (12.0 h day/12.0 h night). In the third experiment, plants (the wall experiment) were also planted in October/2014 but harvested in April/2015 (6 months old) (11.3 h day/12.7 h night). The orientation of the sugarcane was north to south. In the wall experiment, a 2 m high wall made of plywood sheets was built between the first and the second rows on the east side of the field two days before harvest day. The time of harvesting was normalized to a photoperiod of 12 h day/ 12 h night using the following equations to compare the rhythms of samples harvested in different seasons: for times during the daytime, ZT = 12*T*Pd^(−1), where ZT is the normalized time, T is the time from dawn (in hours), and Pd is the length of the day (in hours); for times during the nighttime, ZT = 12 + 12*(T - Pd)*Pn^(−1), where ZT is the normalized time, T is the time from dawn (in hours), Pd is the length of the day (in hours), and Pn is the length of the night (in hours).

### Metabolome

Metabolites were extracted from all 3 biological replicates. First, 50 mg of the grounded tissue was used for MTBE: methanol: water 3:1:1 (v/v/v) extraction, as described previously (Giavalisco *et al*., 2011). Next, the 100 μl of the organic phase was dried and derivatized as previously described (Roessner *et al*., 2001). Finally, 1 μl of the derivatized samples were analyzed on a Combi-PAL autosampler (Agilent Technologies, St. Clara, USA) coupled to an Agilent 7890 gas chromatograph coupled to a Leco Pegasus 2 time-of-flight mass spectrometer (LECO, St. Joseph, MI, USA) in both split (1:40) and splitless modes (Weckwerth *et al*., 2004; Ferreira *et al*., 2018). Chromatograms were exported from Leco ChromaTOF software (version 3.25) to R-3.1.x (R Core Team, 2021). Peak detection, retention time alignment, and library matching were performed using *Target Search R-package* (Cuadros-Inostroza *et al*., 2009). Metabolites were quantified by the peak intensity of a selective mass. Metabolites intensities were normalized by dividing the fresh-weight, followed by the sum of total ion count and global outlier replacement as described previously (Giavalisco *et al*., 2011). Each metabolite value was further normalized to Z-score. The principal component analysis was performed using *pcaMethods* Bioconductor R package. Data ellipses were drawn for the samples harvested during the day and the night (0.85 confidence level).

### Oligoarray hybridizations

Oligoarrays were hybridized as described before (Hotta *et al*., 2013; Dantas *et al*., 2020). Briefly, frozen samples were pulverized in dry, and 100 mg were used for total RNA extraction using Trizol (Life Technologies, Carlsbad, USA). The RNA was treated with DNase I (Life Technologies, Carlsbad, USA) and cleaned using the RNeasy Plant Mini kit (Qiagen, Hilden, Germany) following the supplier’s recommendations. The total RNA quality was assayed using an Agilent RNA 6000 Nano Kit Bioanalyzer chip (Agilent Technologies, St. Clara, USA). Labelling was done following the Low Input Quick Amp Labelling protocol of the Two-Color Microarray-Based Gene Expression Analysis system (Agilent Technologies, St. Clara, USA). Hybridizations were done using a custom 4×44 k oligoarray (Agilent Technologies, St. Clara, USA) previously described (Lembke *et al*., 2012; Hotta *et al*., 2013). Two hybridizations were done for each time point against an equimolar pool of all samples of each organ. Each duplicate was prepared independently using dye swaps. Data were extracted using the Feature Extraction software (Agilent Technologies, St. Clara, USA). Background correction was applied to each dataset. A nonlinear LOWESS normalization was also applied to the datasets to minimize variations due to experimental manipulation. Signals that were distinguishable from the local background signal were taken as an indication that the corresponding transcript was expressed. The GenBank IDs of all sugarcane genes mentioned here can be found in the Supplemental Material (Table S1). The complete dataset can be found at the Gene Expression Omnibus public database under the accession number GSE129543 and GSE171222.

### Data analysis

For further analysis, only expressed transcripts in more than 6 of the 12 time points were considered to be expressed. Identification of rhythmic transcripts was made using an algorithm described in previous work (Dantas *et al*., 2020). Analyses were done in R-3.1.x (R Core Team, 2021). All the time series from expressed transcripts were grouped in co-expressed modules using the R package *weighted correlation network analysis* (*WGCNA*) (Langfelder & Horvath, 2008) with the same parameters as before (Dantas *et al*., 2020). Co-expression modules were considered rhythmic or non-rhythmic using JTK-CYCLE (*P*-value of < 0.75) (Hughes *et al*., 2010). LOESS (locally estimated scatterplot smoothing) regression was used to establish the timing of the peak, or the phase, of each rhythmic time series (Dantas *et al*., 2019). As controls, we also analyzed rhythms using only JTK-Cycle (*P*-value of < 0.05) (Hughes *et al*., 2010) and identified only transcripts with significant gene expression profile differences using the R package *maSigPro* (Nueda *et al*., 2014). Euler diagrams were done using the R package *eulerr*. Code to fully reproduce our analysis is available on GitHub (https://github.com/LabHotta/Microenvironments) and archived on Zenodo (http://doi.org/10.5281/zenodo.4645464).

### RT-qPCR analysis

Frozen leaf samples were pulverized on dry ice using a coffee grinder (Model DCG-20, Cuisinart, Stamford, USA). 100 mg of each pulverized samples were used for total RNA extractions using Trizol (Life Technologies, Carlsbad, USA), according to the manufacturer’s protocol. Total RNA was treated with DNase I (Life Technologies, Carlsbad, USA) and cleaned using the RNeasy Plant Mini Kit (Qiagen). Both RNA quality and concentration were checked using Agilent RNA 6000 Nano Kit Bioanalyzer chip (Agilent Technologies, St. Clara, USA). 5 µg of the purified RNA was used in the reverse transcription reactions using the SuperScript III First-Strand Synthesis System for RT-PCR (Life Technologies Cuisinart). The RT-qPCR reactions were done using Power SYBR Green PCR Master Mix (Applied Biosystems, Waltham, USA), 10× diluted cDNA, and specific primers as previously described (Hotta *et al*., 2013). Reactions were placed in 96-well plates and read with the Fast 7500/7500 Real-Time PCR System (Applied Biosystems, Waltham, USA). Ct determination was performed using the Fast 7500/7500 Real-Time PCR System built-in software (Applied Biosystems, Waltham, USA). The 2^−Δ^*^CT^* method was used to calculate relative expression, using *GLYCERALDEHYDE-3-PHOSPHATE DEHYDROGENASE* (*ScGAPDH*) as a reference gene (Hotta *et al*., 2013). All primers used can be found in the Supplemental Material (Table S2).

## Results

We have measured transcriptional rhythms of leaves, internodes 1&2 and internodes 5 of 9­months-old (9 mo) commercial sugarcane (*Saccharum* hybrid, cultivar SP80 3280) grown in a field in previous work (Dantas *et al*., 2020). We noticed that the circadian clock gene *LATE ELONGATED HYPOCOTYL* (*ScLHY*) peaked hours after dawn, which is unusual, as this gene is light-induced in *Arabidopsis* (Mockler *et al*., 2007). To further investigate this disparity, we measured *ScLHY* rhythms in 4 mo sugarcane leaves that had been harvested from the same field (Fig. 1a). *ScLHY* peaked 0.2 h after dawn in 4 mo sugarcane leaves but peaked 2.7 h after dawn in 9 mo sugarcane leaves (Fig. 1b). We also observed a second *ScLHY* peak that starts increasing between 18.0 h and 20.0 h in 4 mo sugarcane (Fig. 1b). In contrast, *ScTOC1* peaked at 11.0 h and 11.3 h after dawn in 4 mo and 9 mo sugarcane leaves, respectively (Fig. 1b). All plants were harvested from the middle of the field to avoid margin effects.

**Fig. 1.**
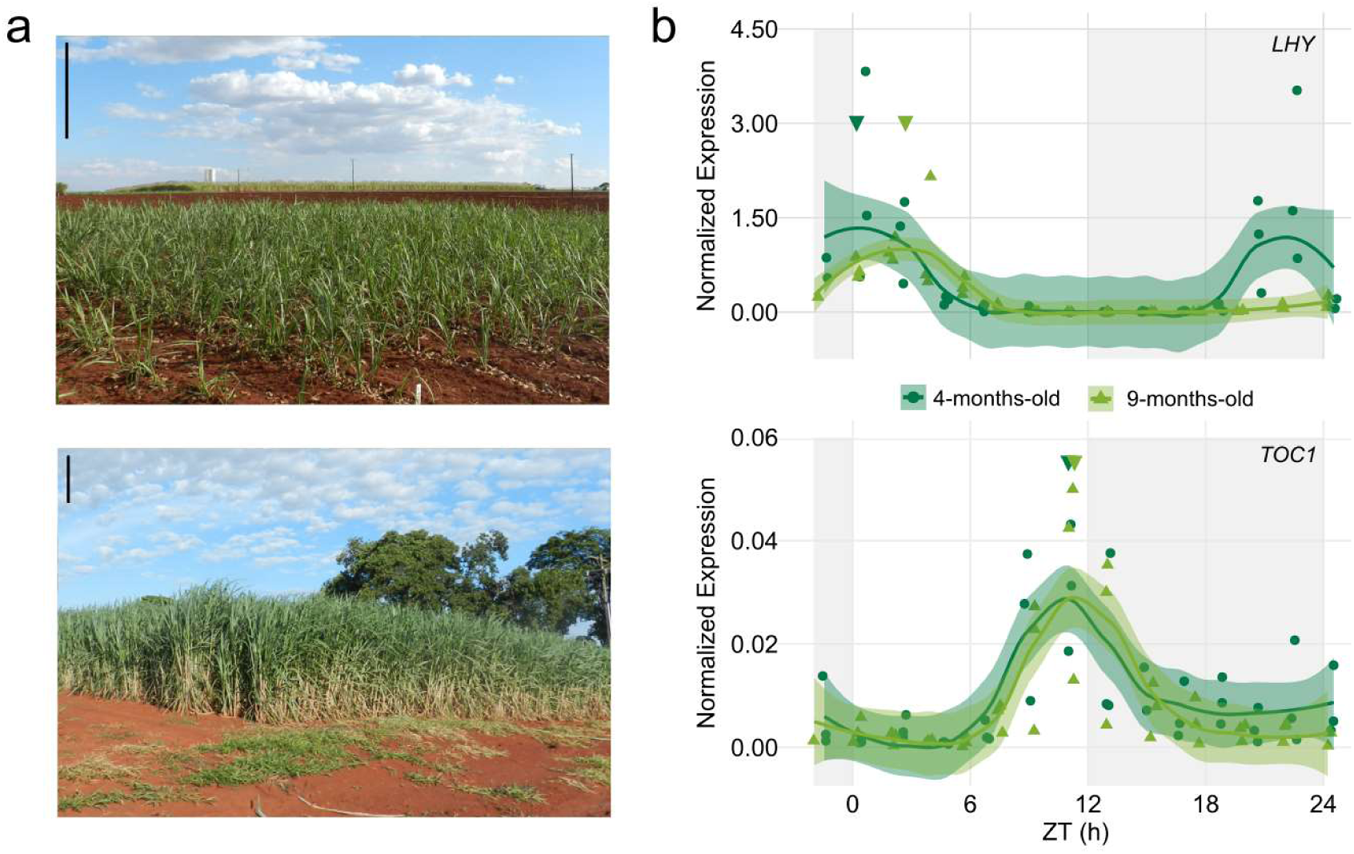
Sugarcane leaves have different *ScLHY* phases in field-grown sugarcane. **(a)** Leaves from sugarcane grown for 4 months old (mo) (upper photo) and 9 mo (lower photo) in the field were harvested for 26 h. Bar = 0.5 m. **(b)** Leaves from sugarcane grown for 4 mo and 9 mo in the field were harvested for 26 h. (A) Diel rhythms of *LATE ELONGATED HYPOCOTYL* (*ScLHY*) and *TIME OF CAB EXPRESSION 1* (*ScTOC1*) in the leaves of 4 mo (dark green) and 9 mo (light green) sugarcane leaves. When sugarcane is 4 months old (left), there is little shading between plants. In comparison, 9 mo sugarcane shade each other when the sun is at a low angle (right). Bar = 0.5 m. All biological replicates and their LOESS curve (continuous lines ± SE) are shown. Inverted triangles show the time of the maximum value of the LOESS curve. Time series were normalized using Z-score. To compare the rhythms of samples harvested in different seasons, the time of harvesting (ZT) was normalized to a photoperiod of 12 h day/ 12 h night. The light-grey boxes represent the night periods. Transcript levels were measured using RT-qPCR. Relative expression was determined using *GLYCERALDEHYDE-3-PHOSPHATE DEHYDROGENASE* (*ScGAPDH*).

To check if other transcriptional rhythms showed a peak change similar to *ScLHY*, we used custom Agilent oligoarrays to measure transcript accumulation in 4 mo sugarcane leaves. We used the transcriptomic data to detect rhythms in 4 mo sugarcane leaves using WGCNA to generate co-expression modules and JTK-Cycle to identify rhythmic modules (Fig. S2). There were 8,553 expressed transcripts in 4 mo sugarcane leaves, 86% of the expressed transcripts found in 9 mo sugarcane leaves (9,891) (Fig. 2a). We considered 4,143 of the expressed transcripts were rhythmic in 4 mo sugarcane leaves (48.4%), lower than in 9 mo sugarcane leaves (68.3%, Fig. S3a). About half of the transcripts that were rhythmic in both 4 mo and 9 mo sugarcane leaves peaked earlier in the older plants (51.5%), whilst 48.2% peaked within 1 h of each other (1.65 ± 1.8 h, median ± MA, n = 4,110) (Fig. 2b). Thus, the phase changes can be seen in transcripts peaking during the whole 24 h cycle, and not just at the same time as *ScLHY* peaks (Fig. S3b). These phase changes were also observed using only JTK-Cycle to identify rhythmic transcripts with phase estimates using JTK_Cycle (1.56 ± 1,9 h, median ± MA, n = 1,014) or using LOESS (3.62 ± 0.4 h, median ± MA, n = 1014) (Figs S4a-d). If only transcripts with significant gene expression profile differences identified with maSigPro were used, the phases differences were 1.65 ± 1.7 h (median ± MA, n = 1,539) using LOESS, and 1.55 ± 2.1 h (median ± MA, n = 527) using JTK-Cycle (Fig. S4e).

**Fig. 2.**
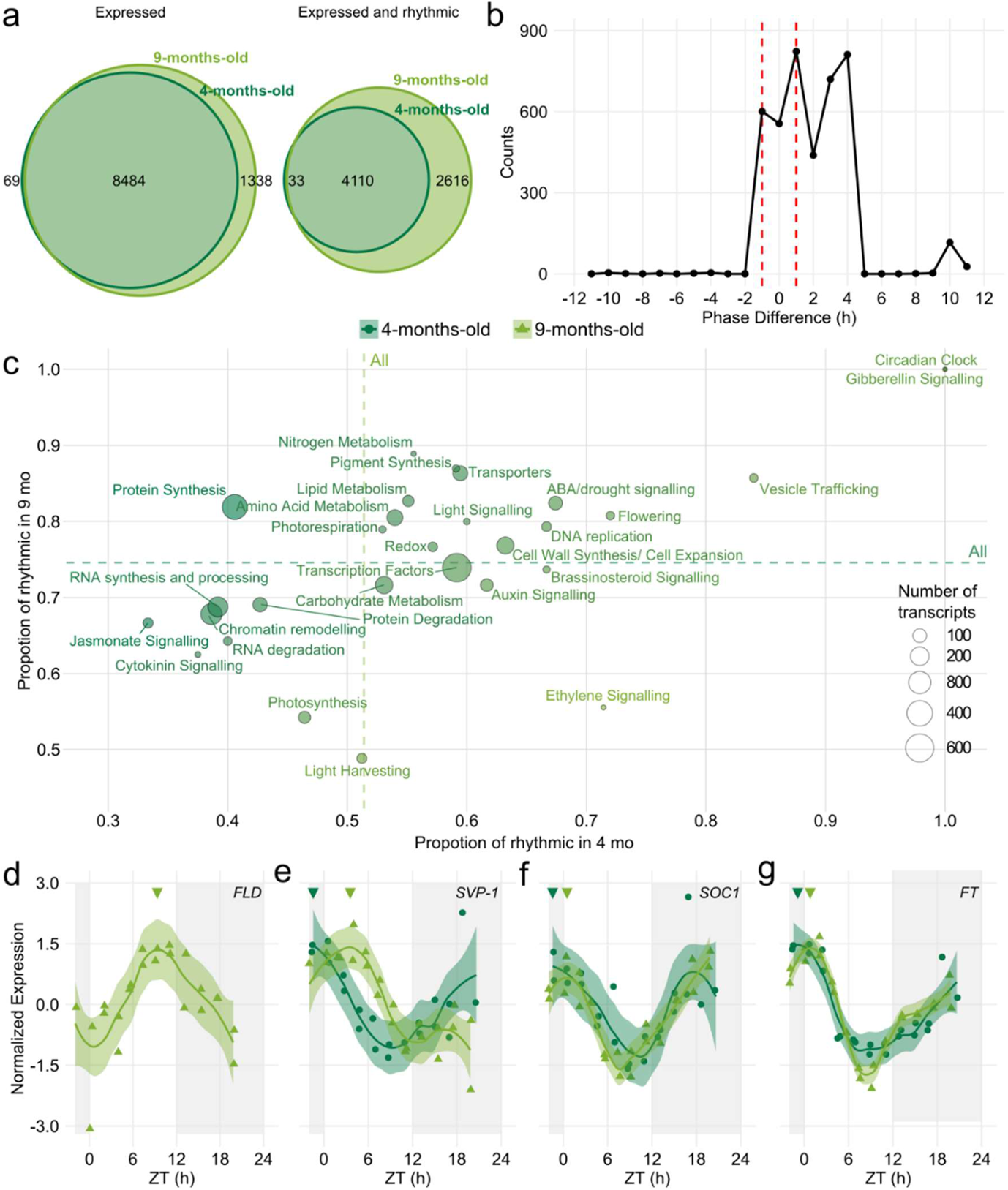
Field-grown sugarcane has transcriptional rhythms with different phases. Leaves from sugarcane grown for 4 months old (4 mo) and 9 mo in the field were harvested for 26 h. **(a)** Euler diagrams of all expressed transcripts and expressed transcripts that are rhythmic in 4 mo (dark green) and 9 mo (light green) sugarcane leaves in diel conditions. **(b)** Distribution of phase differences between rhythmic transcripts from 4 mo and 9 mo sugarcane. Phase differences under ±1 h are not considered significant (red dashed line). **(c)** The proportion of rhythmic transcripts in different functional categories in 4 mo and 9 mo sugarcane leaves. The size of the circles shows the total number of expressed transcripts in each category. The colour of the circles shows if there is an increase in the proportion of rhythmic transcripts in 4 mo (light green) or in 9 mo sugarcane leaves (dark green). The dotted lines represent the proportion of rhythmic transcripts among all annotated transcripts. **(d)** *FLOWERING LOCUS D* (*ScFLD*), **(e)** *SHORT VEGETATIVE-1* (*ScSVP*-1), (**f**) *SUPPRESSOR OF CONSTANS OVEREXPRESSION* (*ScSOC1*), and (**g**) *FLOWERING LOCUS T* (*ScFT*) rhythms measured in 4 mo and 9 mo sugarcane. All biological replicates and their LOESS curve (continuous lines ± SE) are shown. Inverted triangles show the time of the maximum value of the LOESS curve. Time series were normalized using Z-score. To compare the rhythms of samples harvested in different seasons, the time of harvesting (ZT) was normalized to a photoperiod of 12 h day/ 12 h night. The light-grey boxes represent the night periods.

Among the other central oscillator genes investigated, *PSEUDO RESPONSE REGULATOR 73* (*ScPRR73*), *ScPRR59*, *ScGI, REVEILLE 8* (*ScRVE8*) and *EARLY FLOWERING 3* (*ScELF3*) also peaked at similar times in 4 mo and 9 mo sugarcane leaves, while *ScPRR95* had an earlier peak (­2.6 h) in 9 mo sugarcane leaves (Fig. S5). We also have analyzed the proportion of rhythmic transcripts in different functional categories (Figs 2c and S3c). The *Circadian Clock*, *ABA/drought signalling*, *Gibberellin Signalling*, *Transporters*, and *Flowering* are examples of functional categories with a higher proportion of rhythmic transcripts than the average. Conversely, *Chromatin Remodeling* and *RNA Synthesis and processing* are functional categories with lower rhythmic transcripts than the average. Among the disproportionately rhythmic categories in either 4 mo or 9 mo sugarcane leaves, *Auxin Signalling* and *Ethylene Signalling* have a higher proportion of rhythmic transcripts in 4 mo sugarcane leaves. In comparison, *Protein Synthesis* has a higher proportion of rhythmic transcripts in 9 mo sugarcane leaves (Figs 2c and S3c).

One important pathway that could be affected by phase changes in rhythmic genes is flowering signalling. Several sugarcane genes associated with flowering had their transcription patterns changed. For example, *FLOWERING LOCUS D* (*ScFLD*), a histone acetylase, is only expressed in 9 mo sugarcane leaves, with a peak at ZT9 (Fig. 2d). Mutation of *FLD* in Arabidopsis (He *et al*., 2003) and RNA interference of its homolog in rice(Shibaya *et al*., 2016) delays flowering. The flowering genes: *CONSTITUTIVE PHOTOMORPHOGENIC 1* (*ScCOP1*) (Tanaka *et al*., 2011), *CYCLING DOF FACTOR1* (*ScCDF1*)(Higgins *et al*., 2010), *SHORT VEGETATIVE* (*ScSVP-1* and *-3*) (Higgins *et al*., 2010), *SUPPRESSOR OF CONSTANS OVEREXPRESSION 1* (*ScSOC1*) (Higgins *et al*., 2010), *FLOWERING LOCUS T* (Sc*FT*) (Abdul-Awal *et al*., 2020), and *APETALA* (*ScAP1*) (Preston & Kellogg, 2006) peak an average 2 h earlier in 4 mo sugarcane leaves, except for *ScSVP-2* (Figs 2e-f and S6).

We also measured metabolite rhythms in 4 mo and 9 mo sugarcane leaves (Fig. 3). The metabolic profiles of plants harvested during the day and the night cluster separately, except for the leaves harvested 21 h after dawn (zeitgeber time 21, ZT21) in 4 mo sugarcane and leaves harvested at ZT01 in 9 mo sugarcane (Fig. 3a). 31 out of 57 metabolites (54.4%) were found rhythmic in 4 mo sugarcane leaves, whereas 33 out of 51 metabolites (64.7%) were found rhythmic in 9 mo sugarcane leaves. Most of the 20 metabolites that were rhythmic in both plants (75%) peaked later in 9 mo sugarcane (Figs 3b and S7). Thus, the phase changes observed in *ScLHY*, but not in *ScTOC1*, are also widespread among transcriptional and metabolic rhythms. Only 2 metabolites, Sucrose and myo-Inositol, peaked later in 4 mo sugarcane than in 9 mo sugarcane (Figs 3b and S8a). As myo-Inositol has a broader peak, possibly due to a second peak, in 4 mo sugarcane compared to 9 mo sugarcane, this phase-estimate might not be accurate (Fig. S8a). Three metabolites peaked within a 1 h difference (Figs 3d and S8b-c), 10 metabolites peaked 1.5 h to 6 h later in 9 mo sugarcane (Fig. 3e and Fig. S9), and 5 metabolites peaked 8 h to 13 h later in 9 mo sugarcane (Figs 3f and S10). The atypical night peak seen in *ScLHY* was also observed in metabolites such as Alanine (Fig. 3e), Phenylalanine (Fig. S8b), Leucine (Fig. S9b) and Glycerate (Fig. S9e).

**Fig. 3.**
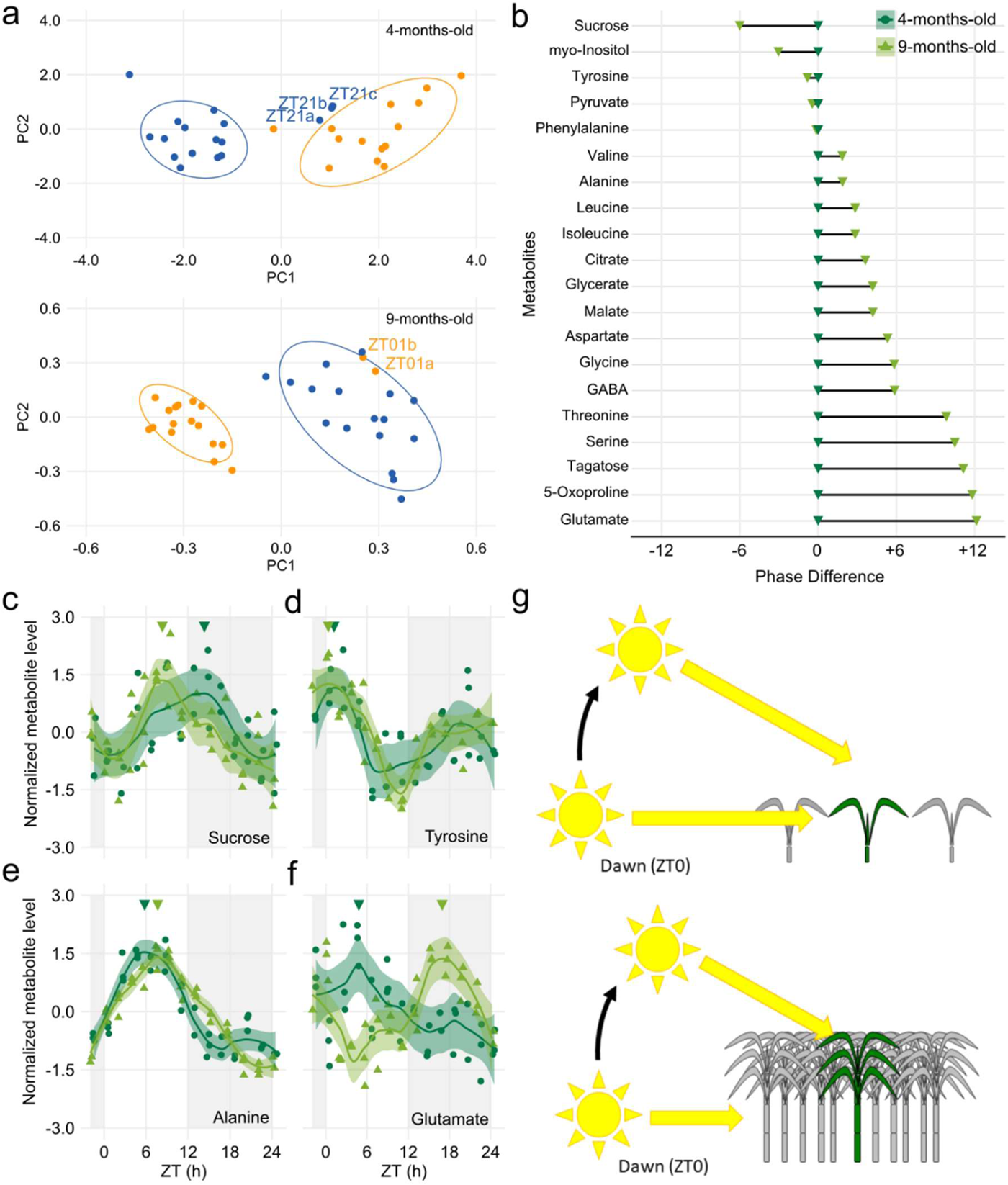
Field-grown sugarcane has metabolic rhythms with different phases. **(a)** Principal Component Analysis (PCA) of the metabolic data from the leaves of 4 mo and 9 mo sugarcane during the day (yellow) or during the night (blue). The percentages of total variance represented by the principal component 1 (PC1) and principal component 2 (PC2) is 63.7% and 60.3%. Metabolites identified in less than 70% of the samples and samples with less than 70% of the total metabolites identified were excluded from the PCA. Data ellipses were drawn for the samples harvested during the day and the night (0.85 confidence level). **(b)** Phase difference in metabolites that were considered rhythmic in both 4 mo (dark green) and 9 mo plants (light green). **(c-f)** Rhythms of **(c)** Sucrose, **(d)** Tyrosine, **(e)** Alanine, and **(f)** Glutamate in 4 mo (dark green) and 9 mo plants (light green). (g) When sugarcane is 4 months old (left), there is little shading between plants. In comparison, 9 mo sugarcane shade each other when the sun is at a low angle (right). All biological replicates (circles in 4 mo or triangles in 9 mo) and their LOESS curve (continuous lines ± SE) are shown. Inverted triangles show the time of the maximum value of the LOESS curve. Metabolite levels were normalized with Z-score. To compare the rhythms of samples harvested in different seasons, the time of harvesting (ZT) was normalized to a photoperiod of 12 h day/ 12 h night. The light-grey boxes represent the night period.

In Arabidopsis, *AtLHY* is responsive to light and temperature, marking the perception of dawn by the plant circadian oscillator(Martínez-García *et al*., 2000; Gould *et al*., 2006). Thus, we hypothesized that the observed phase changes result from the circadian oscillator of 9 mo sugarcane perceiving dawn later than 4 mo sugarcane due to shading effects from neighbouring plants. While young sugarcane plants interact little with their neighbours due to their short height, mature sugarcane plants usually grow densely surrounded by other plants and senescent leaves (Fig. 1a), shading each other, and making it difficult for low angle sunlight to reach leaves uniformly in the entire field (Fig. 3g). This hypothesis also explains why the metabolic profile of 9 mo sugarcane leaves harvested at ZT01, before the *ScLHY* peak, was closer to the leaves harvested during the night than the plants harvested during the day (Fig. 3a).

If our hypothesis were correct, plants that grow on the east side, which receives direct sunlight at dawn, and the west side of a field would have *ScLHY* rhythms with different phases (Fig. 4a). Using leaves from 6 mo plants, we estimated that *ScLHY* from sugarcane peaked 0.6 h after dawn on the east side of the field and 1 h later on the west side (Fig. 4b). Light levels were also around 1 h late on the west side in the morning (Fig. S1c). Moreover, the *ScLHY* transcript levels changed around 800 x during the day in sugarcane on the east side and 350 x in sugarcane on the west side. The peak of ScLHY was 4.5 x higher in sugarcane on the east side, affecting the phase and amplitude of downstream rhythms (Ni *et al*., 2009). *ScPRR73* and *ScPRR37* on the east side show one broad peak, while they have two smaller peaks on the west side (Figs S11a,b). In contrast, *ScTOC1* rhythms had similar phases (Fig. 4b). We concluded that the location of a plant within a field could change the perception of dawn by the circadian clock due to differences in the inner field microenvironment, even if no developmental differences between plants on each side of the field could be detected. In a similar experiment, grape berries (*Vitis vinifera* L.) harvested from the east and the west side of a vine had different phases in sucrose rhythms (Reshef *et al*., 2019).

**Fig. 4.**
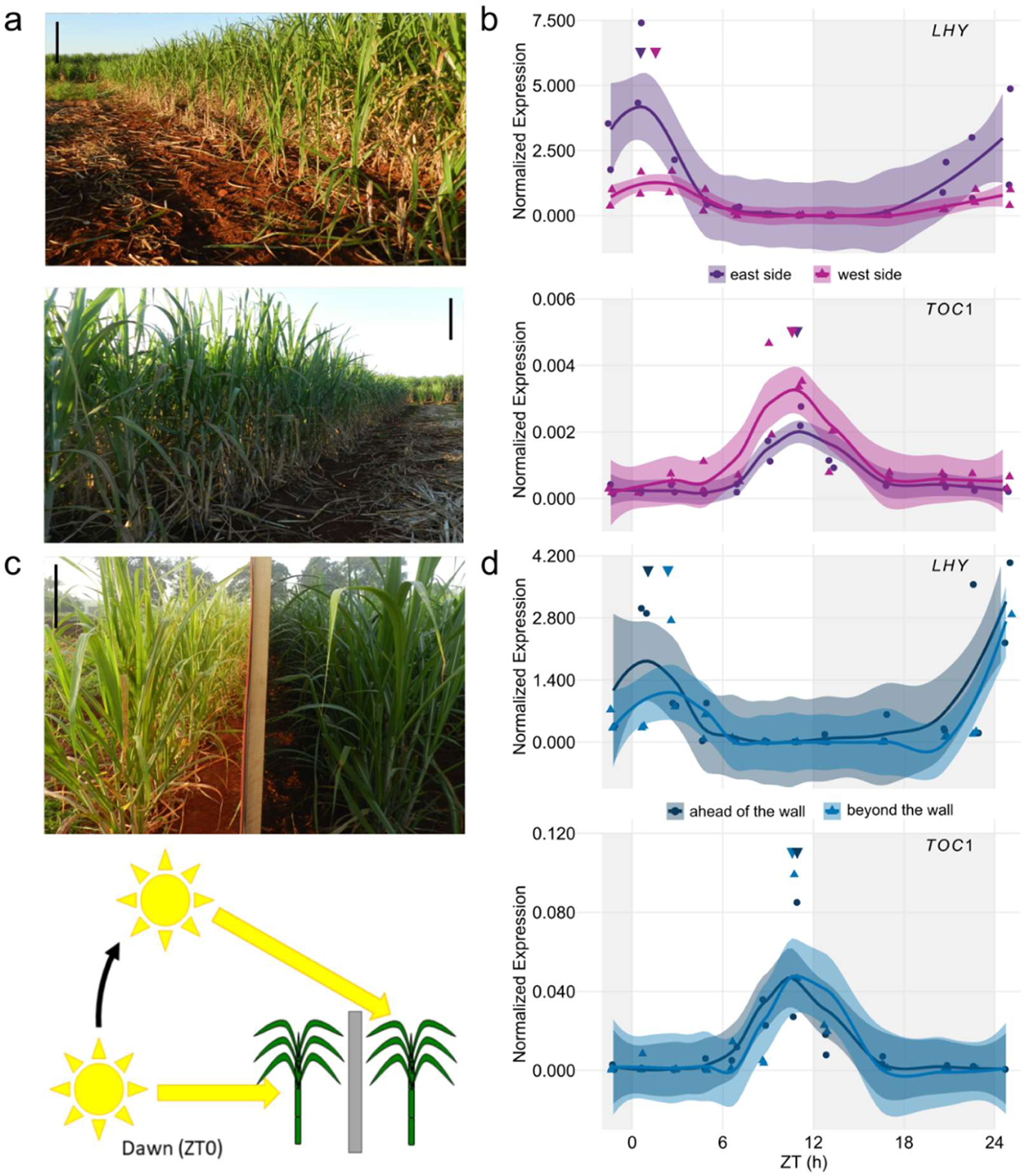
Sugarcane leaves have different *ScLHY* phases when shaded at dawn. **(a)** Leaves from sugarcane grown on the east side (upper photo) and the west side (bottom photo) of the field were harvested for 26 h. Bar = 0.5 m. **(b)** Diel rhythms of *LATE ELONGATED HYPOCOTYL* (*ScLHY*) and *TIME OF CAB EXPRESSION 1* (*ScTOC1*) in the leaves of sugarcane grown on the east side (purple) and the west side (pink) of the field. **(c)** Leaves from sugarcane grown before a wooden wall and after the wooden wall were harvested for 26 h. The photo was taken 1 h after dawn and horizontally flipped. Bar = 0.5 m. **(d)** Diel rhythms of *ScLHY* and *ScTOC1* in sugarcane leaves grown ahead of the wall (dark blue) and beyond the wall (light blue). All biological replicates and their LOESS curve (continuous lines ± SE) are shown. Inverted triangles show the time of the maximum value of the LOESS curve. Time series were normalized using Z-score. To compare the rhythms of samples harvested in different seasons, the time of harvesting (ZT) was normalized to a photoperiod of 12 h day/ 12 h night. The light-grey boxes represent the night periods. Transcript levels were measured using RT-qPCR. Relative expression was determined using *GLYCERALDEHYDE-3-PHOSPHATE DEHYDROGENASE* (*ScGAPDH*).

To further confirm our hypothesis, we created artificial shading by building a wooden wall between the first two lines of 5 mo sugarcane on the east side of the field (Fig. 4c). The wall was built two days before harvesting to avoid developmental effects and any long-term shade avoidance responses caused by shading. Plants ahead the wall received direct sunlight at dawn, while plants beyond the wall received direct sunlight 0.7 h later (Fig. S1e), affecting the local temperature (Fig. S1f). In sugarcane growing ahead of the wall, *ScLHY* expressed in leaves peaked 1.1 h after dawn, whereas *ScLHY* peaked 2.4 h after dawn in leaves beyond the wall (Fig. 4d). The *ScLHY* transcript levels changed 400 x during the day in sugarcane ahead of the wall and 950x in sugarcane beyond the wall. The peak of *ScLHY* was 1.7 x higher in sugarcane ahead of the wall. The *ScPRR73* and *ScPRR37* amplitudes are lower in sugarcane beyond the wall. *ScPRR37* also peaks 1.3 h later in sugarcane beyond the wall (Figs S11c,d). *ScTOC1* peaked at the same time in both conditions (Fig. 4d). These last two experiments show that the phase changes seen in 4 mo and 9 mo sugarcane leaves are not a consequence of seasonal or developmental differences, as plants of the same age that were harvested simultaneously were used. Combining the three experiments, the peaks of *ScLHY* and *ScTOC1* happened 10.4 ± 0.5 h (n = 3) apart in plants not subjected to shading at dawn, and 8.7 ± 0.5 h (n = 3) apart in plants shaded at dawn, which could be enough to trigger photoperiodic responses.

## Discussion

We found that the location of a plant within a field can change the perception of dawn by the circadian clock due to differences in field microenvironments. The circadian clock regulates carbon allocation, which may impact agronomic traits (Yazdanbakhsh *et al*., 2011; Kölling *et al*., 2015). Among the circadian clock genes, only *ScLHY* and *ScPRR95* had their phase changed when comparing 4 mo and 9 mo sugarcane leaves, explaining why only half of the rhythmic genes had phase changes. However, change in the expression of these two genes can trigger changes that affect the timing of a vast number of output rhythms, as hundreds of genes are regulated directly by *AtLHY* and *AtPRR95* in Arabidopsis (Harmer *et al*., 2000; Liu *et al*., 2016; Adams *et al*., 2018).

Diel rhythms could be driven by environmental rhythms, the circadian oscillator, or their synergic combination. Among the external factors, temperature, light quality and quantity, photoperiod and photosynthesis are known to be essential regulators of plant rhythms (Martínez-García *et al*., 2000; Gould *et al*., 2006; Frank *et al*., 2018; Dantas *et al*., 2019). While it is tempting to attribute the phase differences to light, temperature may be an important factor driving sugarcane rhythms. In *Brachypodium dystachion* (L.) P. Beauv., another grass species, changes in temperatures were found to be the main environmental cue driving rhythmic behaviour (Matos *et al*., 2014; MacKinnon *et al*., 2020). In sugarcane, temperature also regulates the rate of *ScLHY* isoform formation by alternative splicing (Dantas *et al*., 2019). Temperature rhythms were very different when 4 mo sugarcane was harvested (temperature between 10°C and 27°C) from when 9 mo sugarcane was harvested (temperature range between 16°C and 30°C) (Fig. S1b). This temperature difference may also explain the differences in the number of rhythmic transcripts and metabolites (Pyl *et al*., 2012; Annunziata *et al*., 2018). In Arabidopsis, differences in temperatures affected the levels of *AtLHY*, *AtCCA1* and *AtPRR9* (a probable ortholog of *ScPRR95*), which included a delay in peak time (Bieniawska *et al*., 2008; Gould *et al*., 2013; Annunziata *et al*., 2018). We should also stress that the 4 mo and 9 mo sugarcane leaves were under different photoperiods (11.5 h day/12.5 h and 13.3 h day/10.7 h night, respectively), which could be another factor explaining some differences among rhythms, as seen in *Arabidopsis halleri* subsp. *gemmifera* (Matsum.) O’Kane & Al-Shehbaz in natural conditions (Nagano *et al*., 2019). However, plants in the other experiments were harvested at the same time.

Sugarcane rhythms may also be regulated by shade avoidance responses, as the R: FR ratio could be altered due to shading from other plants. Even though little is known about the regulation of the circadian clock by shade-avoidance responses, changes in plant density have been correlated to changes in the circadian clock genes. In *Sorghum bicolor* (L.) Moench. at high and low densities, changes in transcription levels of *SbLHY* and *SbTOC1* were correlated with differences in internode length (Kebrom *et al*., 2020). As only one time point was sampled in that study, if sorghum had similar phase differences as the ones we detected, they would be seen as differences in expression levels, as seen when only one timepoint is considered in our experiments (Fig. S12). When row spacings in sugarcane fields were changed between 1.5 m and 2.3 m, cane yields (t/ha) were maintained, as the stalk numbers and weight would change in response to different plant densities (Garside *et al*., 2009; Chiluwal *et al*., 2018).

Microenvironment changes due to shading also affect photosynthesis. In *Arabidopsis*, photosynthetic soluble sugars have been shown to regulate the circadian oscillator (Haydon *et al*., 2013; Frank *et al*., 2018). Hence, the phase changes seen in *ScLHY* could be driven by changes in photosynthesis or, specifically, by carbon assimilation into sucrose. Moreover, photosynthesis is one of the factors that impact rhythms in gene expression in *A. halleri* subsp. *gemmifera* in natural conditions (Nagano *et al*., 2019). It is unclear how the circadian oscillator and metabolites rhythms are interconnected in sugarcane. However, Sucrose was one of the only metabolites with an earlier phase in the leaves of 9 mo sugarcane (Figs 3b,c). Indeed, Sucrose levels in the first 4 h of the day were similar in 4 mo and 9 mo plants. Thus, Sucrose from photosynthesis might not be the factor explaining the observed phase changes, but our analysis only covered a few photosynthetic metabolites. As rhythmic differences in Sucrose were distinct from the changes observed in other metabolites, it is possible that it is dependent on a different internal or external factor. One possible explanation is a reduction of the photosynthesis in the afternoon due to shading of sugarcane in the middle of the field, or the inhibition of photosynthesis due to Sucrose accumulation in the leaves and internodes (McCormick *et al*., 2008).

In 4 mo sugarcane leaves, *ScLHY* and other transcripts showed an unexpected transcriptional peak in the middle of the night in 4 mo sugarcane (Figs 1b and S3a). We do not know what could have triggered this peak in transcript expression, but metabolites, such as Alanine, Glycerate and Phenylalanine, showed the same peak. A similar peak in the middle of the night was found in *Coffea arabica* L. and attributed to moonlight effects (Breitler *et al*., 2020). In our experiment, the increase in transcriptional and metabolic levels during the night started minutes after the crescent moon was set (ZT17-18). Thus, we do not think moonlight or other artificial lights with similar intensity or spectrum were the factors driving these increases. However, this night peak of unknown origin shows that metabolites and transcripts are interconnected and that the metabolic changes brought by photosynthesis and light signalling are not the only force driving rhythms during the day. For example, we have seen significant phase shifts in metabolites (> 8h) between 4 mo and 9 mo sugarcane. The differences, seen in Glutamate, GABA, Serine, 5-Oxoproline, Tagatose and GABA (Figs 3f, S9i and S10), may be due to differences in plant age. In *Nicotiana tabacum* (L.), a similar phase change was observed in the rhythms of Glutamate and GABA in the sink and the source leaves (Masclaux-Daubresse *et al*., 2002). This change was associated with the rhythmic expression of *GLUTAMATE DEHYDROGENASE 1* (*NtGDH1*) in the source leaves but no expression in sink leaves. In 4 mo sugarcane, an *ScGDH1* was arrhythmic, with expression below the detection levels in many time points (Fig. S5i). In 9 mo sugarcane, *ScGDH1* was rhythmic, and expression was above detection levels in all time points.

Flowering is a necessary process that could be affected by phase changes in rhythmic genes. As the vegetative stage is harvested, flowering is usually undesired in sugarcane, as it reduces sucrose yield due to decreased growth, increased side-shoot production, and pithiness (Rao, 1977). Sugarcane enters the reproductive phase in response to the shortening of the photoperiod over 15 days, and just a 0.5 h difference in photoperiod can trigger flowering in most genotypes (Midmore, 1980; Moore & Berding, 2013; Glassop *et al*., 2014). Thus, the correct measurement of dawn and dusk is essential for flowering in sugarcane. Among those genes, *ScSVP*-1, *ScSVP*-3, *ScSOC1*, *ScFT*, and *ScCDF1* peak before dawn in 4 mo sugarcane leaves and after dawn in 9 mo sugarcane leaves (Figs 2e-g and S6). In Arabidopsis, AtLHY and AtCCA1 reduce AtSVP protein levels, a repressor of flowering (Fujiwara *et al*., 2008). A change in the relative phase between *ScLHY* and *ScSVP* may change the ScSVP protein levels, impacting flowering. We also found that *ScPRR37* and *ScPRR73* show different expression patterns in plants shaded in the morning that could be explained by the changes in the phase of two peaks during the day. In sorghum, the second peak in *SbPRR37*, during the evening, regulates flowering (Murphy *et al*., 2011). Thus, neighbour shading could trigger flowering earlier than the actual critical photoperiod. In sugarcane, the spindle, a whorl of immature leaves on the top of the plant, perceives photoperiod, which could be a strategy to reduce the effects of shading from neighbour plants (Moore & Berding, 2013; Glassop *et al*., 2014; Glassop & Rae, 2019).

The circadian clock increases the productivity of plants when its phase matches the phase of environmental rhythms (Dodd *et al*., 2005). It is usually accepted that the rhythms of plants under the same photoperiod have matching phases. However, we have evidence that field microenvironmental rhythms and not astronomical rhythms regulate the plant circadian oscillator of plants in natural, fluctuating conditions and transcriptional and metabolic rhythms. This phenomenon can probably be observed in plants grown in natural or artificial environments, such as crops, forests, and gardens, with microenvironments. Thus, rhythms in field microenvironments are another factor to consider when translating knowledge from the lab to the field, especially agriculture.

## Acknowledgements

This work was supported by the São Paulo Research Foundation (FAPESP) (grant nos. 11/00818­8, 15/06260-0 and 19/08534-0; BIOEN Program) and by the Serrapilheira Institute (grant no. Serra-1708-16001). LLBD and NOL were supported by FAPESP scholarships (grants 11/08897-4 and 16/06740-4, respectively). We thank the support of Carolina G. Lembke for the oligoarrays hybridization. We thank the support of the LabMET at the Brazilian Bioethanol Science and Technology Laboratory (CTBE/CNPEM) for the metabolite profiling analysis (MET-19154 and MET-20673).

## Author Contributions

Conceptualization, LLBD, MSC, CTH; Methodology, LLBD, CTH; Software, CTH, MYN; Validation, LLBD, NOL; Investigation, LLBD, NOL, CTH, CC; Resources, MSC, C. C.; Data Curation, CTH, MYN; Writing – Original Draft, LLBD, CTH; Writing – Review & Editing, LLBD, MSC, CC, CTH; Visualization; C. T. H.; Project Administration, CTH; Funding Acquisition, CTH.

## Data availability

The complete dataset can be found at the Gene Expression Omnibus public database under the accession number GSE129543 and GSE171222. Code to fully reproduce our analysis is available on GitHub (https://github.com/LabHotta/Microenvironments) and archived on Zenodo (http://doi.org/10.5281/zenodo.4645464).

---

**Fig. S1.**
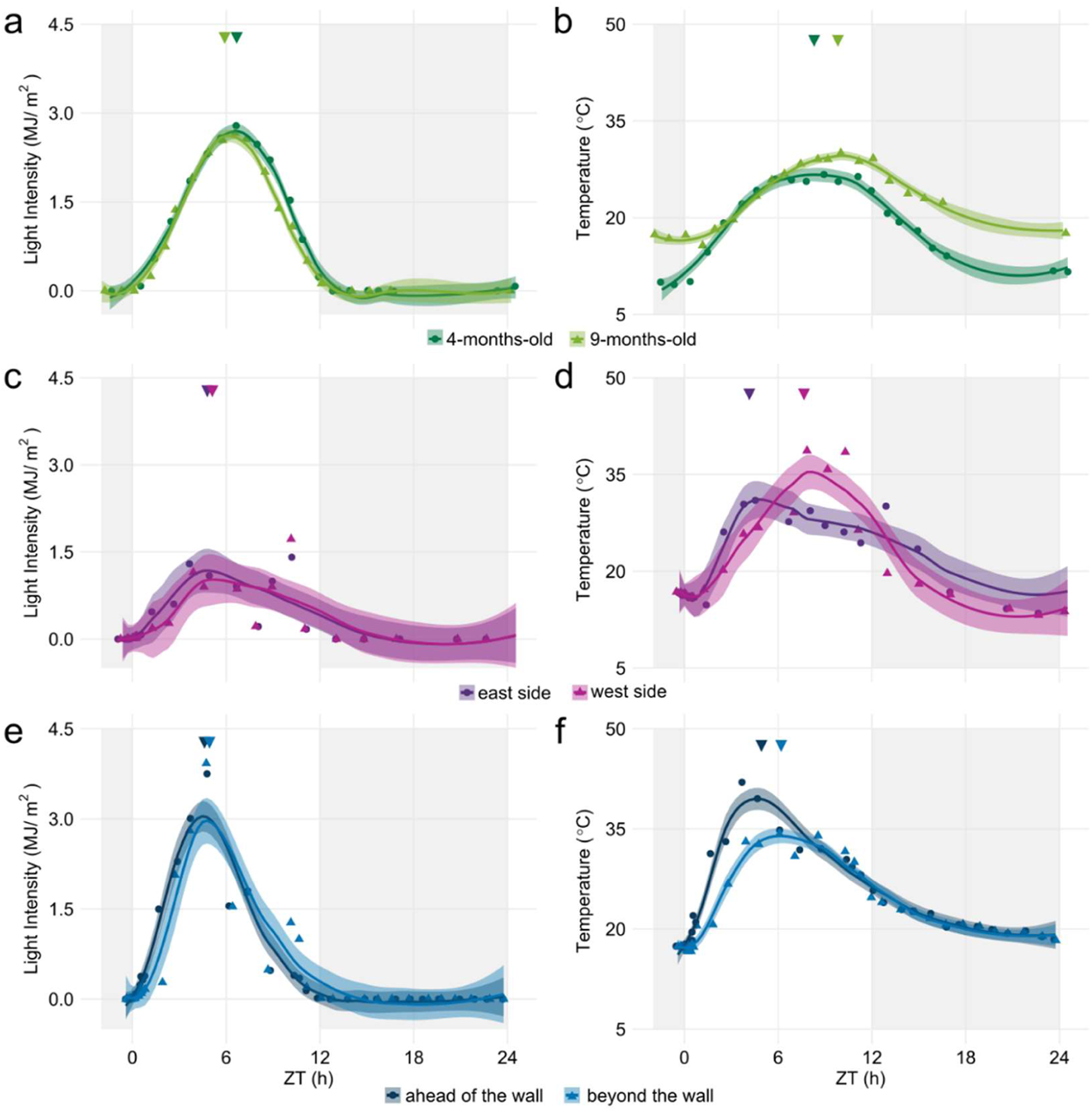
Environmental conditions in Araras (SP, Brazil). **(a)** Light intensity and **(b)** temperature on the day of harvesting 4 months old (4 mo) (green) and 9 mo (light green) sugarcane. Data was taken from a meteorological station nearby the sugarcane field, which underestimates microenvironmental changes. **(c)** Light intensity and **(d)** temperature on the day of harvesting 6 mo sugarcane in the east (purple) or the west side (pink) of the field. Data was taken from a handheld light meter and thermometer next to the plants inside the field. **(e)** Light intensity and **(f)** temperature on the day of harvesting 5 mo sugarcane in the east side of the field (dark blue) and sugarcane shaded by an artificial wall (light blue). Data was taken from a handheld light meter and thermometer next to the plants inside the field. All data and their LOESS curve (continuous lines ± SE) are shown. Inverted triangles show the time of the maximum value of the LOESS curve. To compare the rhythms of samples harvested in different seasons, the time of harvesting (ZT) was normalized to a photoperiod of 12 h day/ 12 h night. The light-grey boxes represent the night period.

**Fig. S2.**
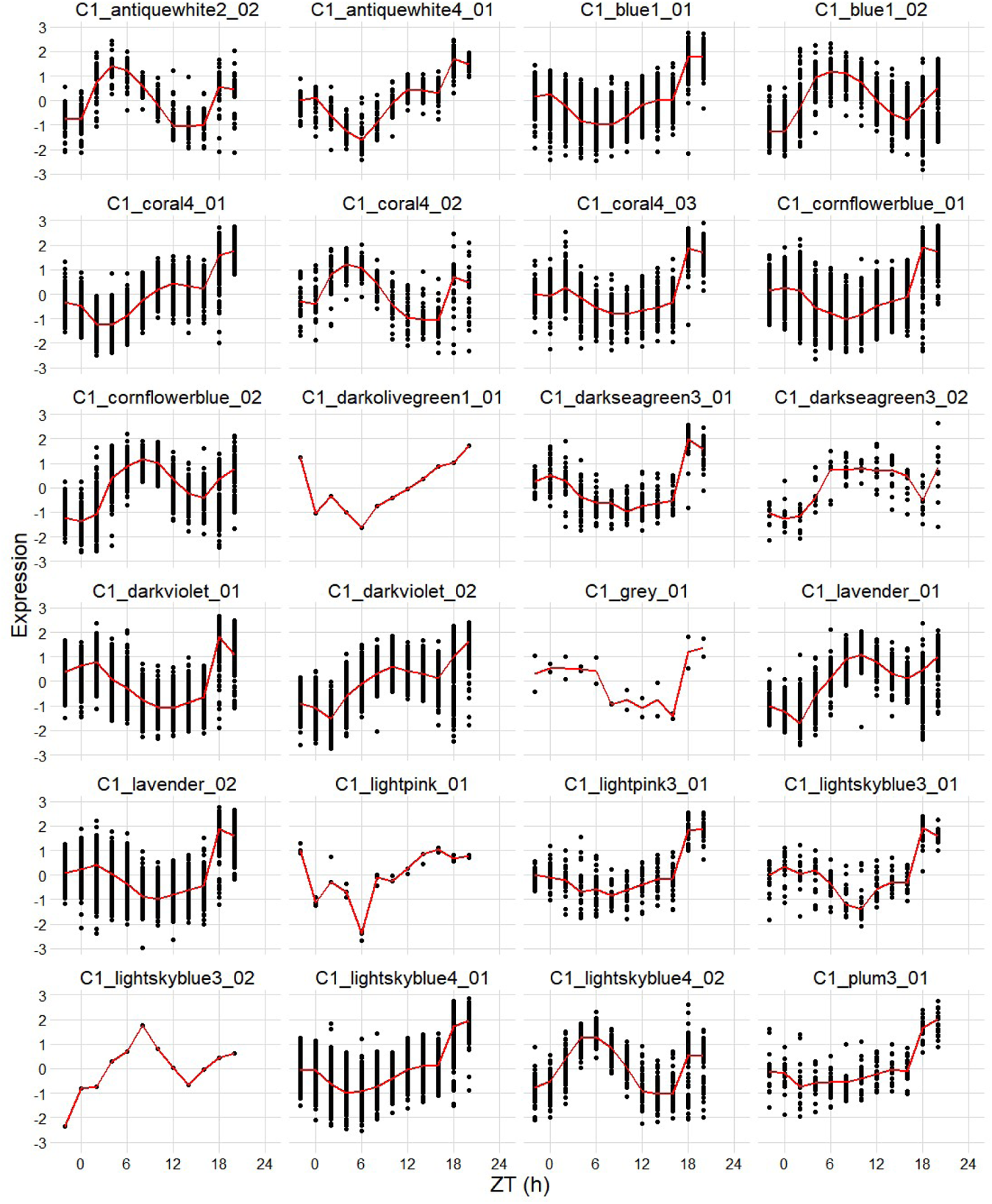
Rhythmic co-expression modules that were identified in 4-months-old sugarcane leaves. Weighted correlation network analysis was used to group co-expressed transcripts in modules. Rhythmic modules were identified using JTK-cycle. The time course generated from the median of all time points is in red.

**Fig. S3.**
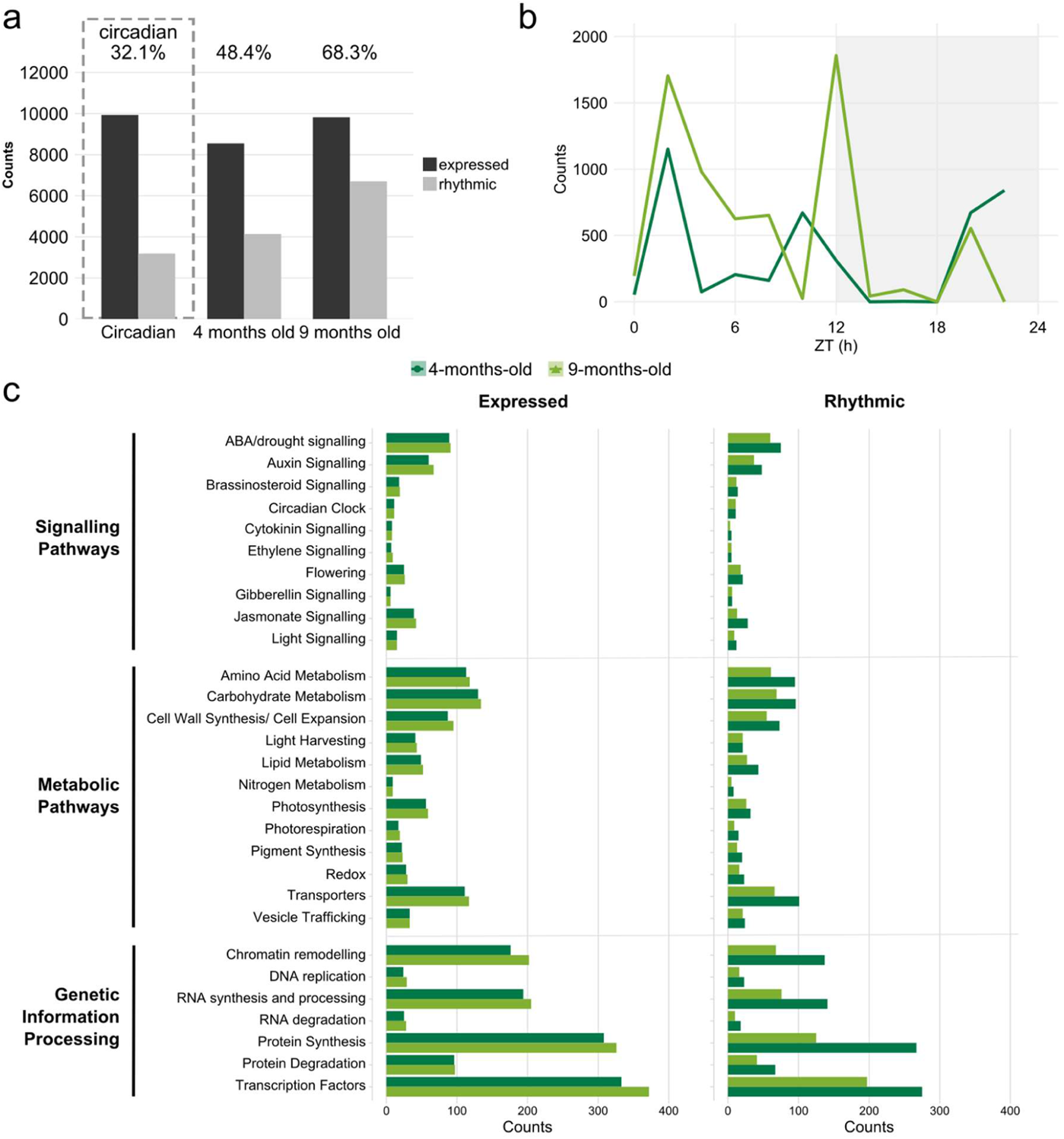
Rhythmic co-expression modules that were identified in 4-months-old sugarcane leaves. **(a)** The number of expressed and rhythmic transcripts detected in the leaves of 4­months-old (4 mo) and 9 mo sugarcane in field-grown (diel) conditions and 3 mo leaves in circadian conditions published in Hotta et al. (2013)(Hotta *et al*., 2013). **(b)** Distribution of the peak time of rhythmic transcripts in 4 mo and 9 mo sugarcane. **(c)** Distribution of the functional categories of expressed and rhythmic transcripts in 4 mo (dark green) and 9 mo (light green) sugarcane.

**Fig. S4.**
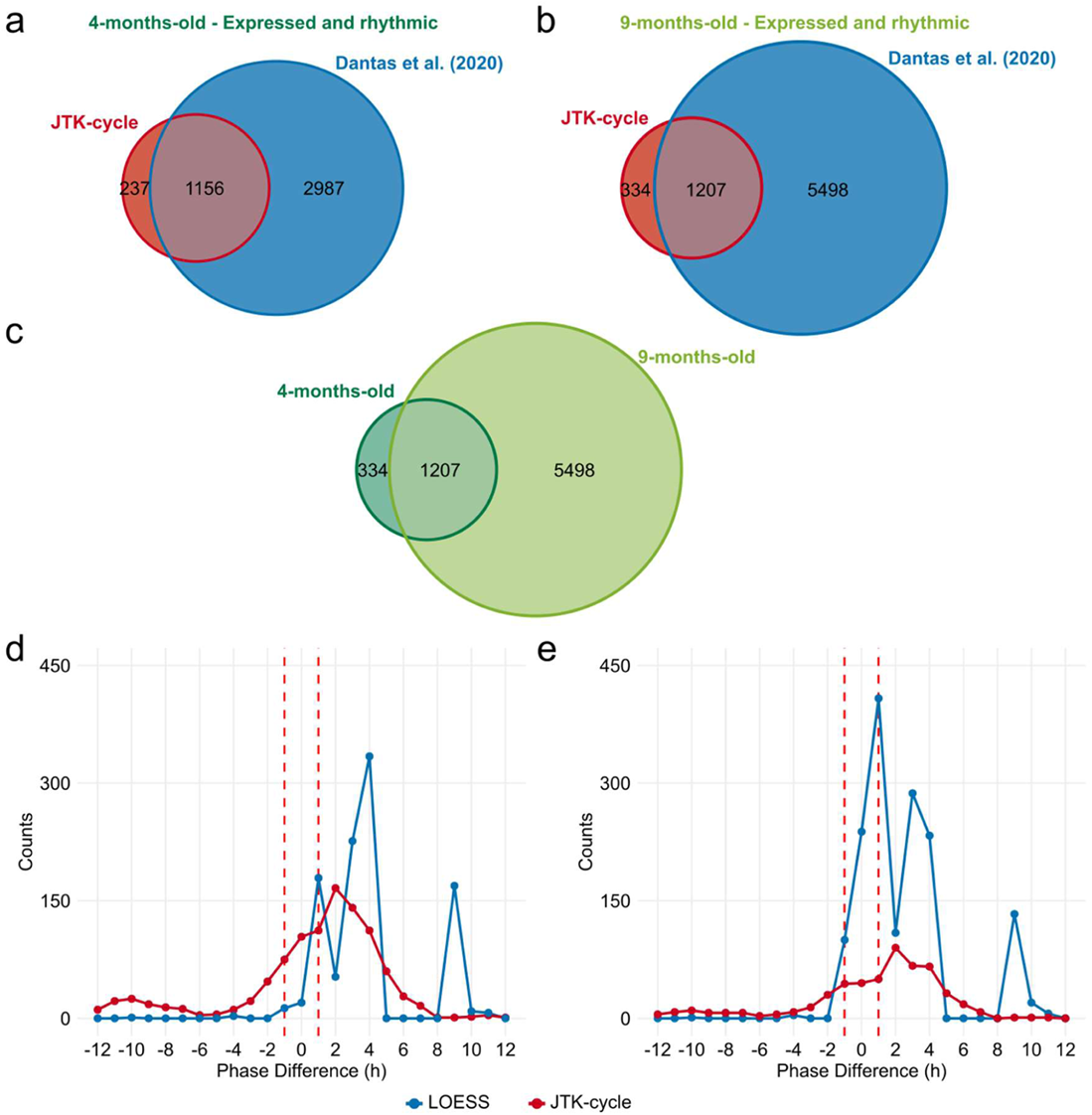
Phase changes between rhythmic transcripts in 4-month-old and 9-month-old sugarcane leaves. **(a-b)** Rhythmic transcripts were identified using JTK-Cycle (blue) and compared with the WGCNA and JTK-Cycle used in Dantas et al. (2020) transcripts (red) in **(a)** 4­month-old (4 mo) and **(b)** 9 mo sugarcane leaves using Euler diagrams. **(c)** Euler diagram of the rhythmic transcripts identified by JTK-Cycle in 4 mo (dark green) and 9 mo sugarcane leaves (green). **(c-d)** Distribution of phase differences between rhythmic transcripts from 4 mo and 9 mo sugarcane using **(d)** JTK-Cycle and **(e)** maSigPro. Phase differences were estimated using JTK-Cycle (red) or LOESS (blue). Differences under ±1 h are not considered significant (red dashed line).

**Fig. S5.**
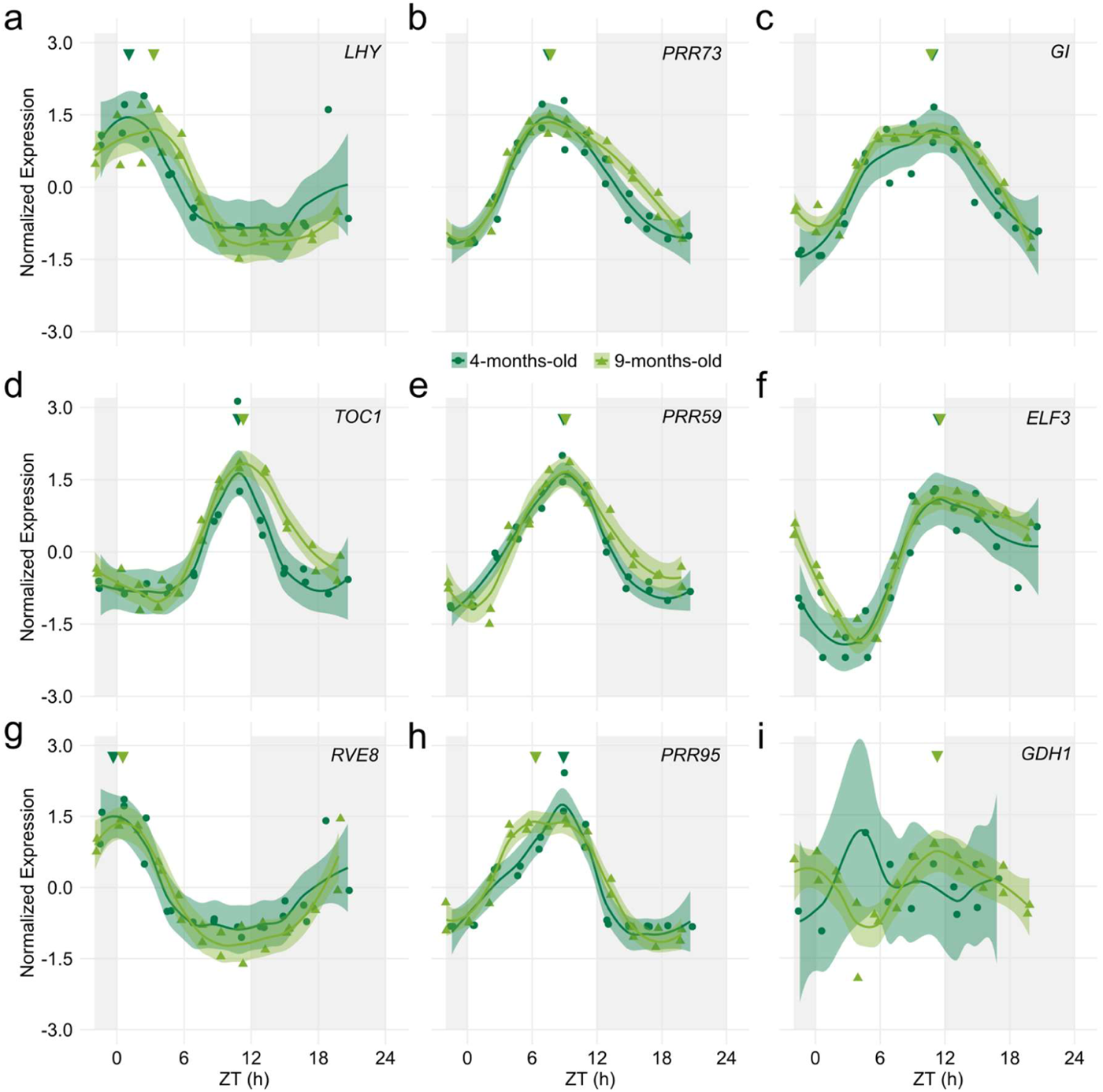
Rhythms of circadian clock genes and N metabolism genes in the leaves of 4 months old and 9 months old sugarcane. Rhythms were measured using custom Agilent oligo arrays in the leaves of 4 mo and 9 mo sugarcane grown in the field. All biological replicates and their LOESS curve (continuous lines ± SE) are shown. Inverted triangles show the time of the maximum value of the LOESS curve. Time series were normalized using Z-score. To compare the rhythms of samples harvested in different seasons, the time of harvesting (ZT) was normalized to a photoperiod of 12 h day/ 12 h night. The light-grey boxes represent the night periods.

**Fig. S6.**
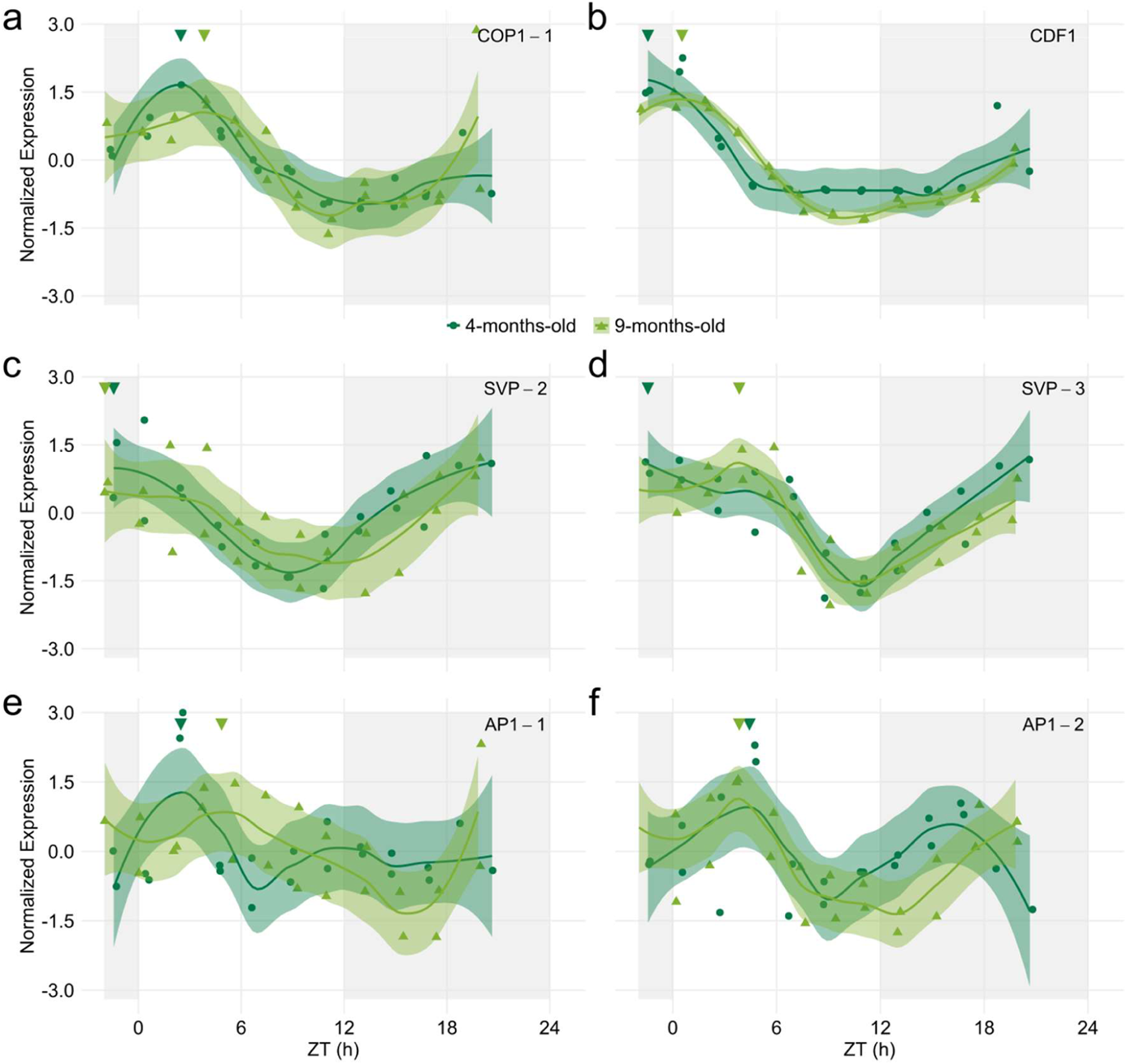
Rhythms of genes associated with flowering in the leaves of 4 months old and 9 months old sugarcane. Rhythms of **(a)** *CONSTITUTIVE PHOTOMORPHOGENIC 1* (*ScCOP1*), **(b)** *CYCLING DOF FACTOR1* (*ScCDF1*), **(c)** *SHORT VEGETATIVE-1* (*ScSVP*-2), **(d)** *ScSVP*-3, **(e)** *APETALA1*-1 (*ScAP1*-1), and **(f)** *ScAP1*-2 were measured using custom Agilent oligo arrays in the leaves of 4 mo and 9 mo sugarcane grown in the field. All biological replicates and their LOESS curve (continuous lines ± SE) are shown. Inverted triangles show the time of the maximum value of the LOESS curve. Time series were normalized using Z-score. To compare the rhythms of samples harvested in different seasons, the time of harvesting (ZT) was normalized to a photoperiod of 12 h day/ 12 h night. The light-grey boxes represent the night periods.

**Fig. S7.**
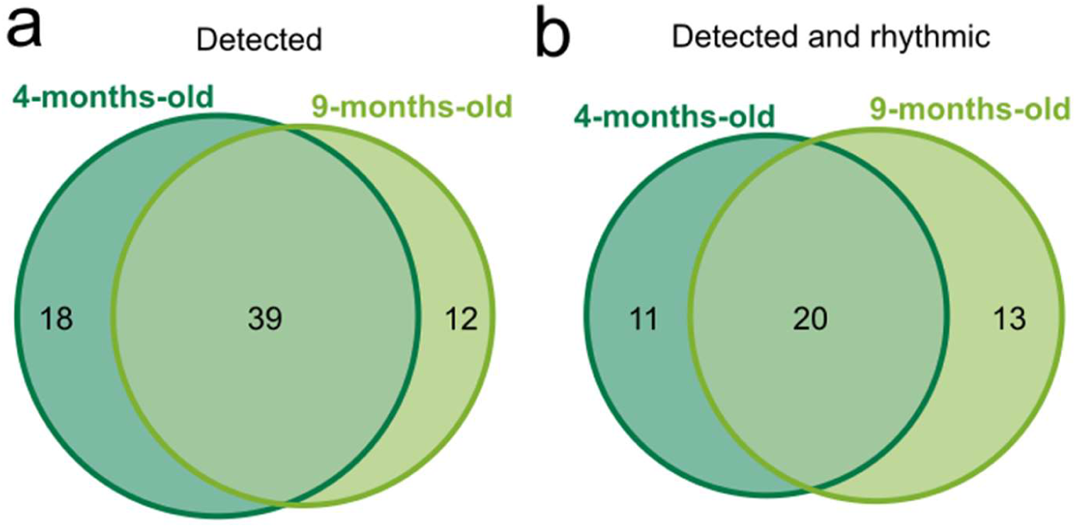
Euler diagram of detected and rhythmic metabolites. Euler diagrams of all **(a)** detected metabolites and **(b)** rhythmic metabolites in 4 mo (dark green) and 9 mo (light green) sugarcane leaves in diel conditions.

**Fig. S8.**
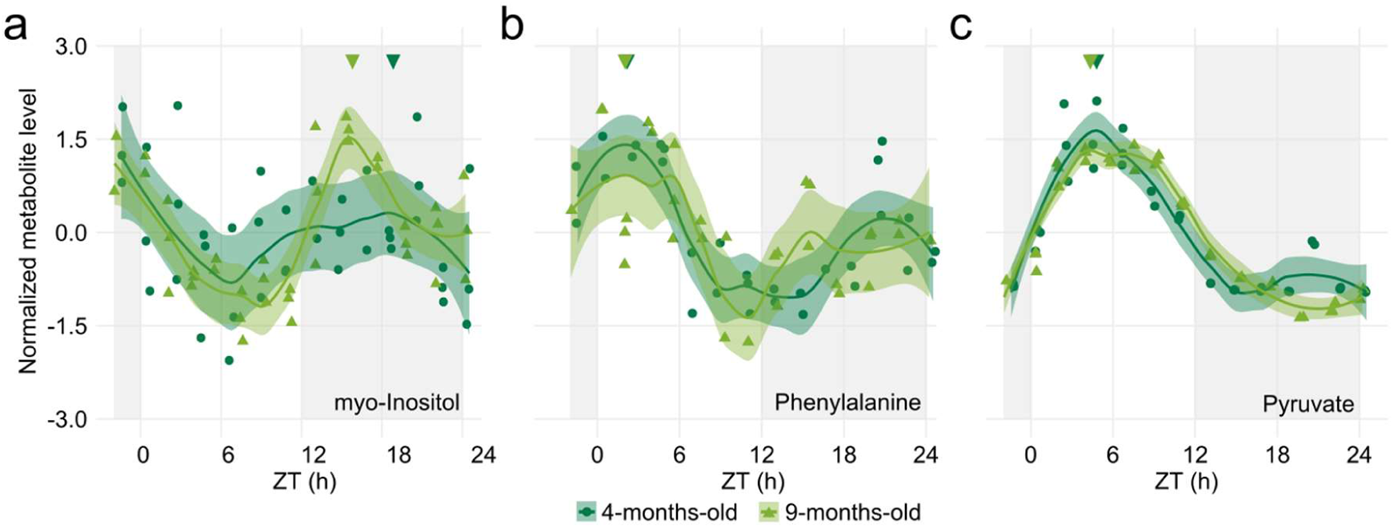
Metabolite rhythms that had an early phase or no phase change in 9 months old sugarcane compared to 4 months old sugarcane. Leaf +1 of sugarcane grown for 4 months old (4 mo, dark green) and 9 mo (light green) in the field were harvested for 26 h. Rhythms of **(a)** myo-Inositol, **(b)** Phenylalanine and **(c)** Pyruvate in 4 mo and 9 mo Metabolite levels were normalized with Z-score. All biological replicates (circles in 4 mo and triangles in 9 mo) and their LOESS curve (continuous lines ± SE) are shown. Inverted triangles show the time of the maximum value of the LOESS curve. To compare the rhythms of samples harvested in different seasons, the time of harvesting (ZT) was normalized to a photoperiod of 12 h day/ 12 h night. The light-grey boxes represent the night period.

**Fig. S9.**
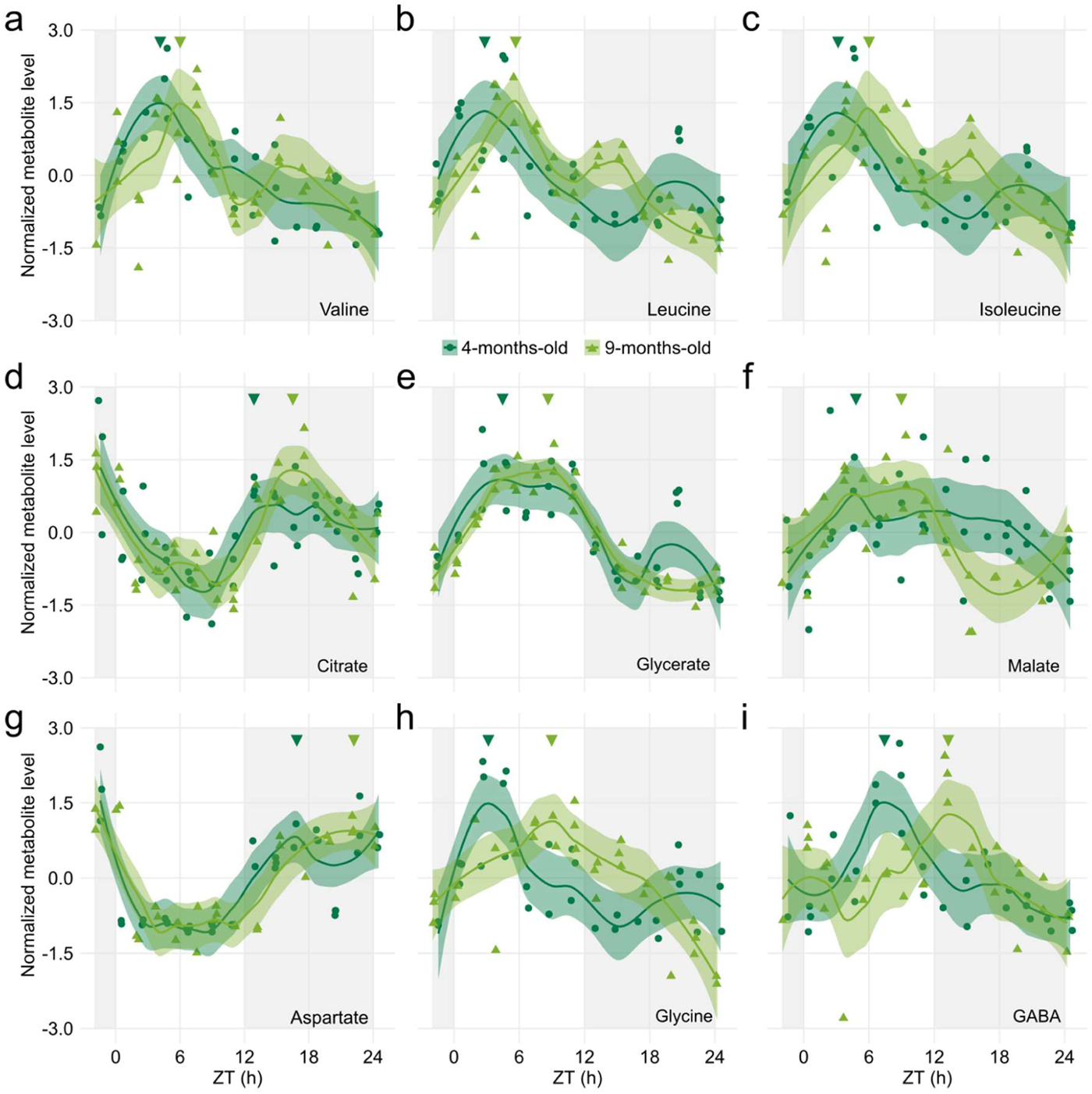
Metabolite rhythms that had a phase increase smaller or equivalent to 6 h in 9 months old sugarcane compared to 4 months old sugarcane. Leaf +1 of sugarcane grown for 4 months old (4 mo, dark green) and 9 mo (light green) in the field were harvested for 26 h. Rhythms of **(a)** Valine, **(b)** Leucine, **(c)** Isoleucine, **(d)** Citrate, **(e)** Glycerate, **(f)** Malate, **(g)** Aspartate, **(h)** Glycine, and **(i)** GABA in 4 mo and 9 mo sugarcane leaves. Metabolite levels were normalized with Z-score. All biological replicates (circles in 4 mo and triangles in 9 mo) and their LOESS curve (continuous lines ± SE) are shown. Inverted triangles show the time of the maximum value of the LOESS curve. To compare the rhythms of samples harvested in different seasons, the time of harvesting (ZT) was normalized to a photoperiod of 12 h day/ 12 h night. The light-grey boxes represent the night period.

**Fig. S10.**
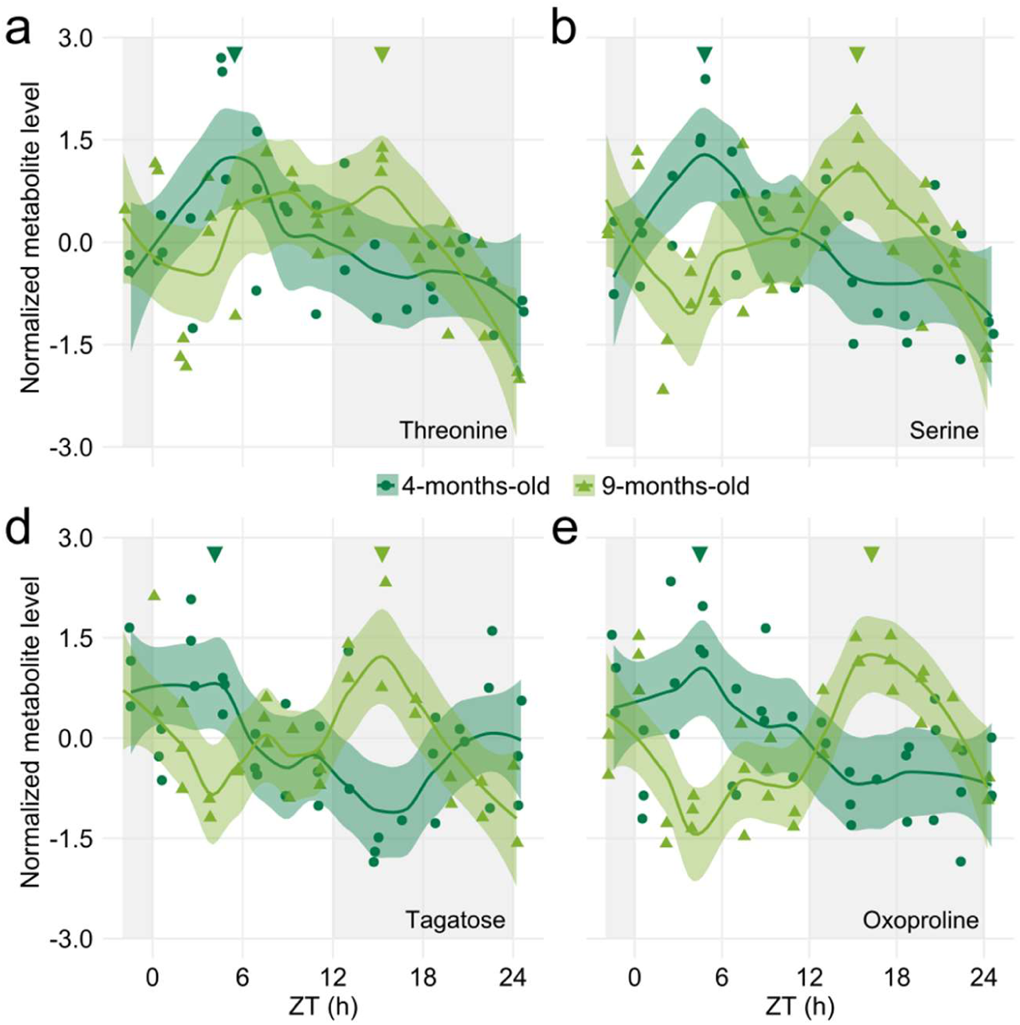
Metabolite rhythms that had a phase increase larger than 6 h in 9 months old sugarcane compared to 4 months old sugarcane. Leaf +1 of sugarcane grown for 4 months old (4 mo, dark green) and 9 mo (light green) in the field were harvested for 26 h. Rhythms of **(a)** Threonine, **(b)** Serine, **(c)** Tagatose and **(d)** Oxoproline in 4 mo and 9 mo Metabolite levels were normalized with Z-score. All biological replicates (circles in 4 mo and triangles in 9 mo) and their LOESS curve (continuous lines ± SE) are shown. Inverted triangles show the time of the maximum value of the LOESS curve. To compare the rhythms of samples harvested in different seasons, the time of harvesting (ZT) was normalized to a photoperiod of 12 h day/ 12 h night. The light-grey boxes represent the night period.

**Fig. S11.**
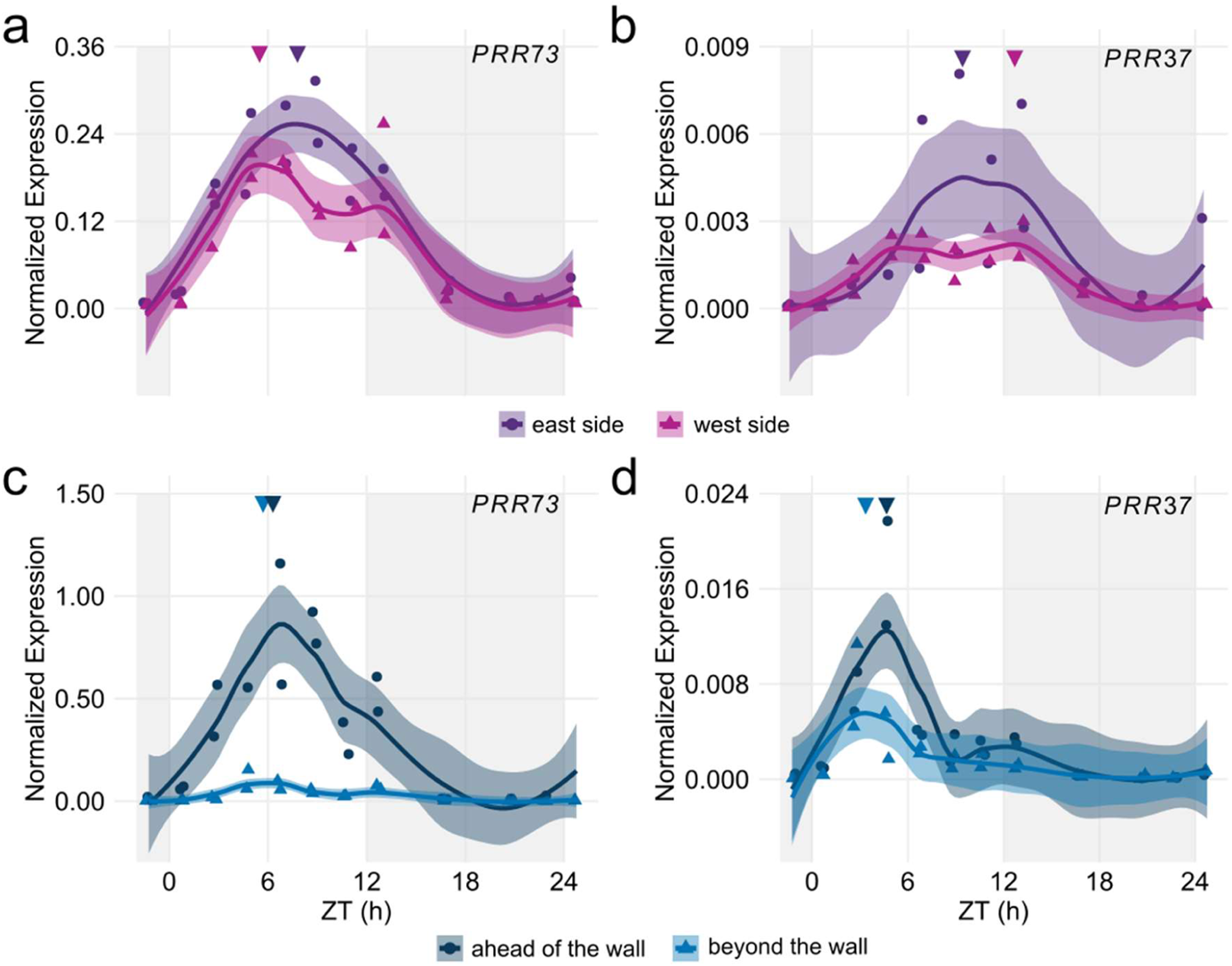
Sugarcane leaves have different *ScPRR73* and *ScPRR37* patterns when shaded at dawn. **(a-b)** Diel rhythms of **(a)** *PSEUDO-RESPONSE REGULATOR 73* (*ScPRR73*) and **(b)** *ScPRR37* in the leaves of sugarcane grown on the east side (purple) and the west side (pink) of the field. **(c-d)** Diel rhythms of **(c)** *ScPRR73* and **(d)** *ScPRR37* in the leaves of sugarcane grown ahead of the wall (dark blue) and beyond the wall (light blue). All biological replicates and their LOESS curve (continuous lines ± SE) are shown. Inverted triangles show the time of the maximum value of the LOESS curve. Time series were normalized using Z-score. To compare the rhythms of samples harvested in different seasons, the time of harvesting (ZT) was normalized to a photoperiod of 12 h day/ 12 h night. The light-grey boxes represent the night periods. Transcript levels were measured using RT-qPCR.

**Fig. S12.**
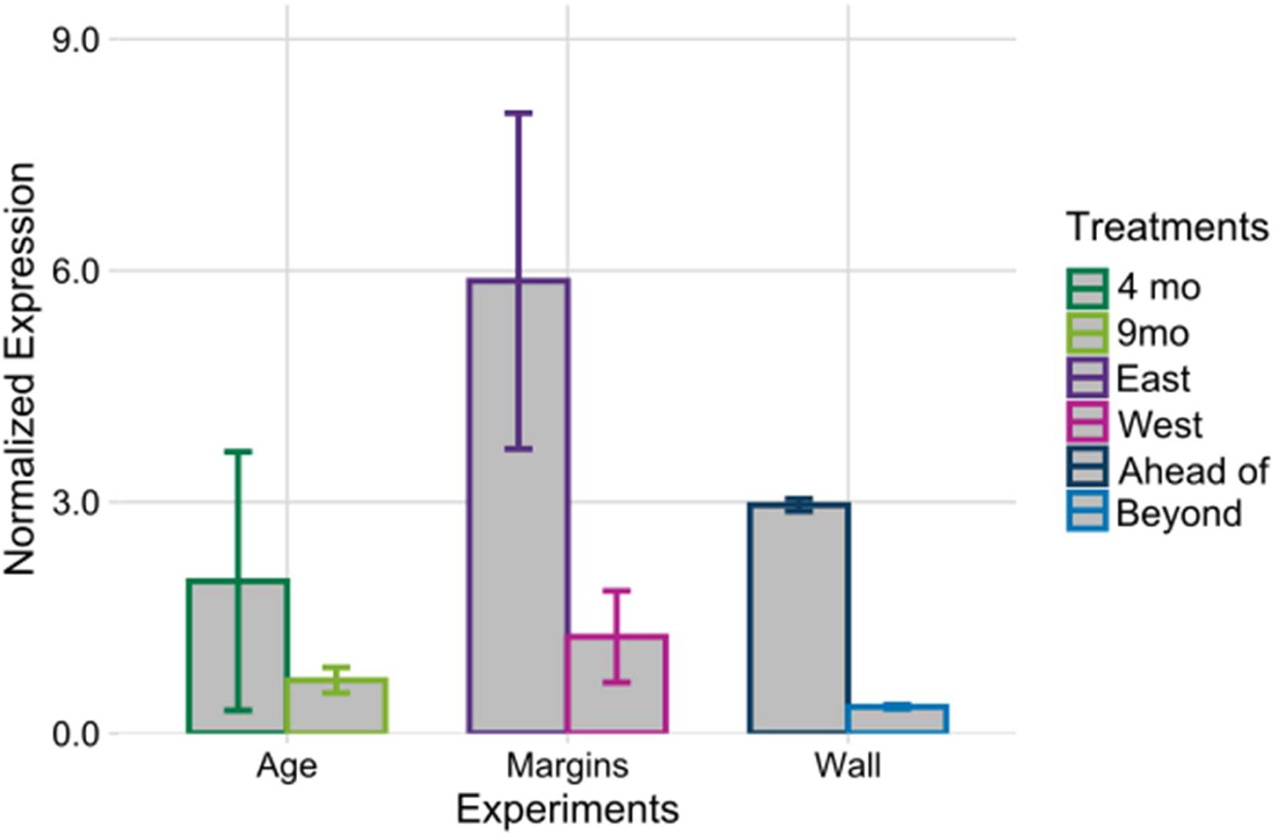
Transcript levels of *ScLHY* in the first hours of the morning. In the Age experiment, 4-months-old (dark green) and 9-months-old (light leaves) sugarcane leaves were used. In the Margins experiment, sugarcane leaves from the East side (purple) or the West side (pink) of the sugarcane field were taken. In the Wall experiment, sugarcane leaves were taken from plants ahead of the wall (dark blue) and beyond the wall (light blue) on the east side of the field. Sugarcane leaves were harvested between ZT0 and ZT2 (n=3). Transcript levels were measured using RT-qPCR. Relative expression was determined using *GLYCERALDEHYDE-3­PHOSPHATE DEHYDROGENASE* (*ScGAPDH*).

**Table S1.**
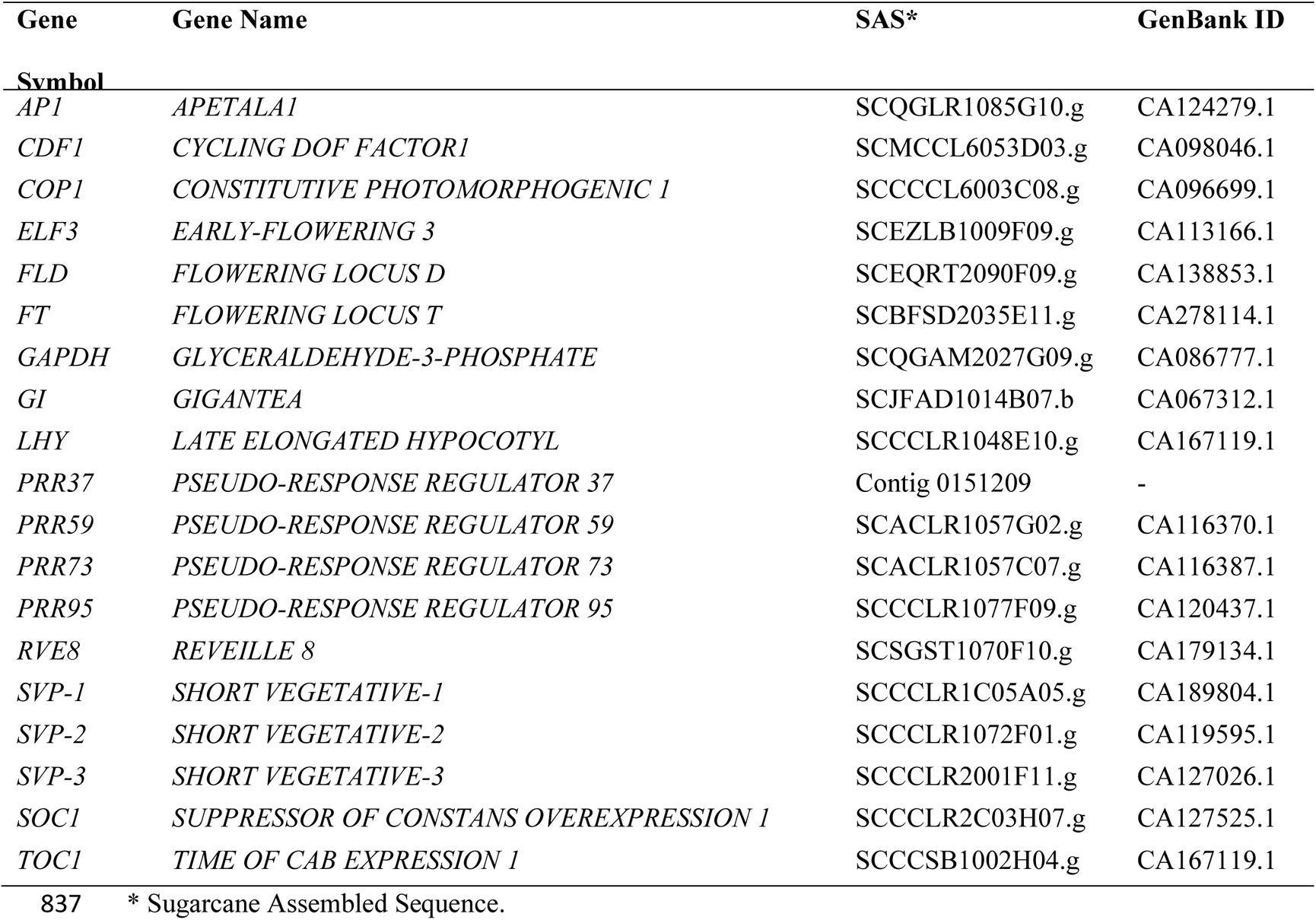
Sugarcane genes and GenBank IDs.

**Table S2.**
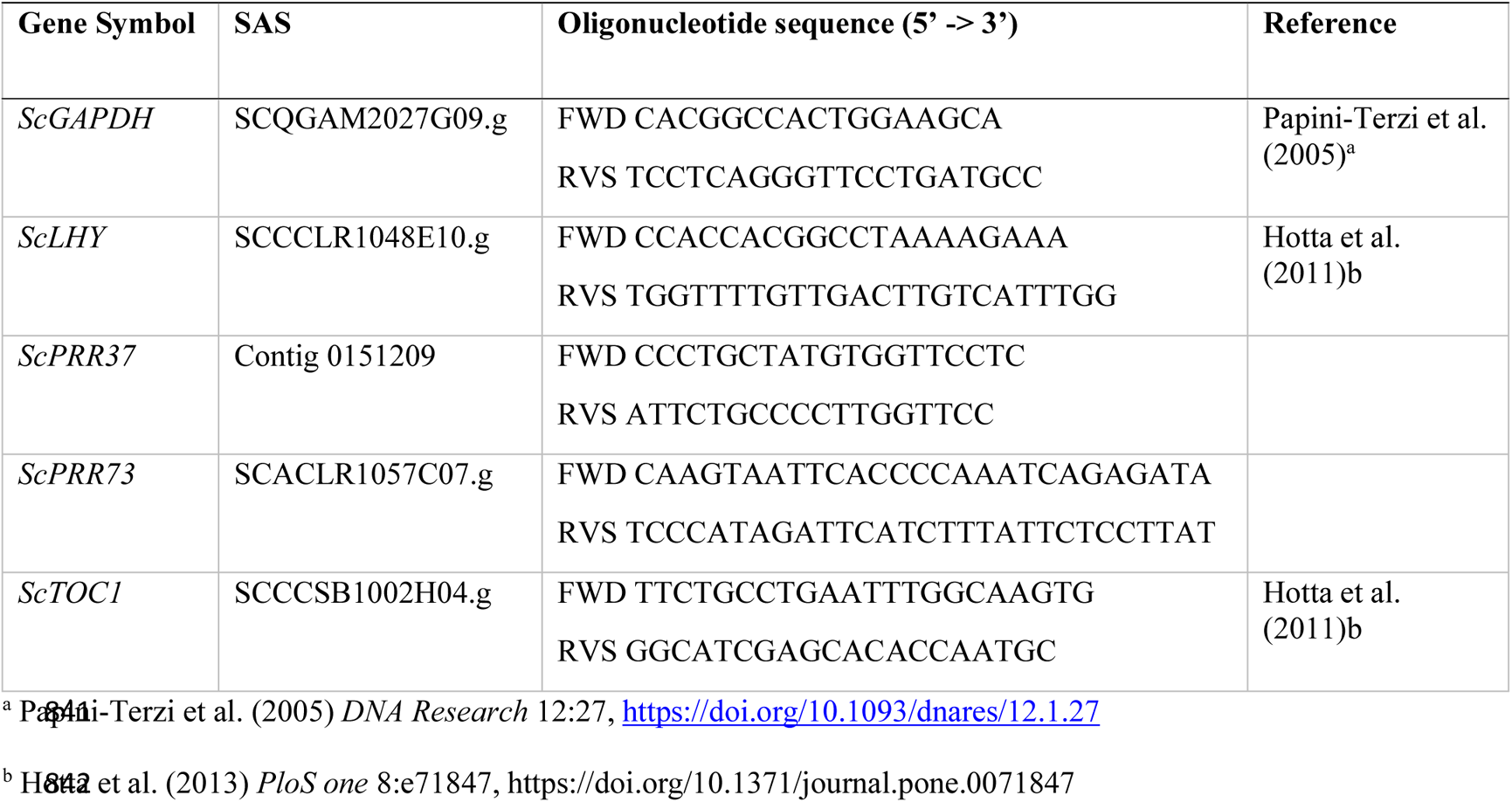
Sugarcane primers pairs used to validate oligo arrays expression levels using RT-qPCR.

## References

1. Abdul-Awal SM, Chen J, Xin Z, Harmon FG. 2020. A sorghum gigantea mutant attenuates florigen gene expression and delays flowering time. Plant Direct 4: e00281.

2. Adams S, Grundy J, Veflingstad SR, Dyer NP, Hannah MA, Ott S, Carré IA. 2018. Circadian control of abscisic acid biosynthesis and signalling pathways revealed by genome-wide analysis of LHY binding targets. New Phytologist 220: 893–907.

3. Alabadí D, Oyama T, Yanovsky MJ, Harmon FG, Más P, Kay SA. 2001. Reciprocal regulation between TOC1 and LHY/CCA1 within the Arabidopsis circadian clock. Science (New York, N.Y.) 293: 880–883.

4. Annunziata MG, Apelt F, Carillo P, Krause U, Feil R, Koehl K, Lunn JE, Stitt M. 2018. Response of Arabidopsis primary metabolism and circadian clock to low night temperature in a natural light environment. Journal of Experimental Botany 69: 4881–4895.

5. Annunziata MG, Apelt F, Carillo P, Krause U, Feil R, Mengin V, Lauxmann MA, Köhl K, Nikoloski Z, Stitt M, et al. 2017. Getting back to nature: a reality check for experiments in controlled environments. Journal of Experimental Botany 68: 4463–4477.

6. Bieniawska Z, Espinoza C, Schlereth A, Sulpice R, Hincha DK, Hannah MA. 2008. Disruption of the Arabidopsis Circadian Clock Is Responsible for Extensive Variation in the Cold-Responsive Transcriptome. Plant Physiology 147: 263–279.

7. Breitler J-C, Djerrab D, Leran S, Toniutti L, Guittin C, Severac D, Pratlong M, Dereeper A, Etienne H, Bertrand B. 2020. Full moonlight-induced circadian clock entrainment in Coffea arabica. BMC Plant Biology 20: 24.

8. Calixto CPG, Waugh R, Brown JWS. 2015. Evolutionary relationships among barley and Arabidopsis core circadian clock and clock-associated genes. Journal of Molecular Evolution 80: 108–119.

9. Cha J-Y, Kim J, Kim T-S, Zeng Q, Wang L, Lee SY, Kim W-Y, Somers DE. 2017. GIGANTEA is a co-chaperone which facilitates maturation of ZEITLUPE in the Arabidopsis circadian clock. Nature Communications 8: 3.

10. Chiluwal A, Singh HP, Sainju U, Khanal B, Whitehead WF, Singh BP. 2018. Spacing Effect on Energy Cane Growth, Physiology, and Biomass Yield. Crop Science 58: 1371–1384.

11. Cuadros-Inostroza A, Caldana C, Redestig H, Kusano M, Lisec J, Peña-Cortés H, Willmitzer L, Hannah MA. 2009. TargetSearch--a Bioconductor package for the efficient preprocessing of GC-MS metabolite profiling data. BMC bioinformatics 10: 428.

12. Dantas LL de B, Almeida-Jesus FM, de Lima NO, Alves-Lima C, Nishiyama-Jr MY, Carneiro MS, Souza GM, Hotta CT. 2020. Rhythms of Transcription in Field-Grown Sugarcane Are Highly Organ Specific. Scientific Reports 10: 6565.

13. Dantas LLB, Calixto CPG, Dourado MM, Carneiro MS, Brown JWS, Hotta CT. 2019. Alternative Splicing of Circadian Clock Genes Correlates With Temperature in Field-Grown Sugarcane. Frontiers in Plant Science 10.

14. Dodd AN, Salathia N, Hall A, Kevei E, Toth R, Nagy F, Hibberd JM, Millar AJ, Webb AA. 2005. Plant circadian clocks increase photosynthesis, growth, survival, and competitive advantage. Science 309: 630–3.

15. Ferreira DA, Martins MCM, Cheavegatti-Gianotto A, Carneiro MS, Amadeu RR, Aricetti JA, Wolf LD, Hoffmann HP, de Abreu LGF, Caldana C. 2018. Metabolite Profiles of Sugarcane Culm Reveal the Relationship Among Metabolism and Axillary Bud Outgrowth in Genetically Related Sugarcane Commercial Cultivars. Frontiers in Plant Science 9.

16. Frank A, Matiolli CC, Viana AJC, Hearn TJ, Kusakina J, Belbin FE, Wells Newman D, Yochikawa A, Cano-Ramirez DL, Chembath A, et al. 2018. Circadian entrainment in Arabidopsis by the sugar-responsive transcription factor bZIP63. Current Biology 28: 2597–2606.e6.

17. Fujiwara S, Oda A, Yoshida R, Niinuma K, Miyata K, Tomozoe Y, Tajima T, Nakagawa M, Hayashi K, Coupland G, et al. 2008. Circadian Clock Proteins LHY and CCA1 Regulate SVP Protein Accumulation to Control Flowering in Arabidopsis. The Plant Cell 20: 2960–2971.

18. Garside AL, Bell MJ, Robotham BG, Garside AL, Bell MJ, Robotham BG. 2009. Row spacing and planting density effects on the growth and yield of sugarcane. 2. Strategies for the adoption of controlled traffic. Crop and Pasture Science 60: 544–554.

19. Giavalisco P, Li Y, Matthes A, Eckhardt A, Hubberten H-M, Hesse H, Segu S, Hummel J, Köhl K, Willmitzer L. 2011. Elemental formula annotation of polar and lipophilic metabolites using (13) C, (15) N and (34) S isotope labelling, in combination with high-resolution mass spectrometry. The Plant Journal: For Cell and Molecular Biology 68: 364–376.

20. Glassop D, Rae AL. 2019. Expression of sugarcane genes associated with perception of photoperiod and floral induction reveals cycling over a 24-hour period. Functional Plant Biology 46: 314–327.

21. Glassop D, Rae AL, Bonnett GD. 2014. Sugarcane Flowering Genes and Pathways in Relation to Vegetative Regression. Sugar Tech 16: 235–240.

22. Gould PD, Locke JCW, Larue C, Southern MM, Davis SJ, Hanano S, Moyle R, Milich R, Putterill J, Millar AJ, et al. 2006. The Molecular Basis of Temperature Compensation in the Arabidopsis Circadian Clock. The Plant Cell 18: 1177–1187.

23. Gould PD, Ugarte N, Domijan M, Costa M, Foreman J, Macgregor D, Rose K, Griffiths J, Millar AJ, Finkenstädt B, et al. 2013. Network balance via CRY signalling controls the Arabidopsis circadian clock over ambient temperatures. Molecular Systems Biology 9: 650.

24. Gray JA, Shalit-Kaneh A, Chu DN, Hsu PY, Harmer SL. 2017. The REVEILLE Clock Genes Inhibit Growth of Juvenile and Adult Plants by Control of Cell Size. Plant Physiology 173: 2308– 2322.

25. Green RM, Tingay S, Wang Z-Y, Tobin EM. 2002. Circadian rhythms confer a higher level of fitness to Arabidopsis plants. Plant Physiology 129: 576–584.

26. Harmer SL, Hogenesch JB, Straume M, Chang HS, Han B, Zhu T, Wang X, Kreps JA, Kay SA. 2000. Orchestrated transcription of key pathways in Arabidopsis by the circadian clock. *Science (New York*, N.Y.) 290: 2110–2113.

27. Haydon MJ, Mielczarek O, Robertson FC, Hubbard KE, Webb AA. 2013. Photosynthetic entrainment of the Arabidopsis thaliana circadian clock. Nature 502: 689–92.

28. He Y, Michaels SD, Amasino RM. 2003. Regulation of flowering time by histone acetylation in Arabidopsis. *Science (New York*, N.Y.) 302: 1751–1754.

29. Herrero E, Kolmos E, Bujdoso N, Yuan Y, Wang M, Berns MC, Uhlworm H, Coupland G, Saini R, Jaskolski M, et al. 2012. EARLY FLOWERING4 recruitment of EARLY FLOWERING3 in the nucleus sustains the Arabidopsis circadian clock. The Plant Cell 24: 428–443.

30. Higgins JA, Bailey PC, Laurie DA. 2010. Comparative Genomics of Flowering Time Pathways Using Brachypodium distachyon as a Model for the Temperate Grasses. PLOS ONE 5: e10065.

31. Hotta CT, Nishiyama MY, Souza GM. 2013. Circadian rhythms of sense and antisense transcription in sugarcane, a highly polyploid crop. PLoS One 8: e71847.

32. Huang H, Gehan MA, Huss SE, Alvarez S, Lizarraga C, Gruebbling EL, Gierer J, Naldrett MJ, Bindbeutel RK, Evans BS, et al. 2017. Cross-species complementation reveals conserved functions for EARLY FLOWERING 3 between monocots and dicots. Plant Direct 1: e00018.

33. Huang W, Pérez-García P, Pokhilko A, Millar AJ, Antoshechkin I, Riechmann JL, Mas P. 2012. Mapping the Core of the Arabidopsis Circadian Clock Defines the Network Structure of the Oscillator. Science 336: 75–79.

34. Hughes ME, Hogenesch JB, Kornacker K. 2010. JTK_CYCLE: an efficient nonparametric algorithm for detecting rhythmic components in genome-scale data sets. Journal of Biological Rhythms 25: 372–380.

35. Izawa T, Mihara M, Suzuki Y, Gupta M, Itoh H, Nagano AJ, Motoyama R, Sawada Y, Yano M, Hirai MY, et al. 2011. Os-GIGANTEA confers robust diurnal rhythms on the global transcriptome of rice in the field. The Plant Cell 23: 1741–1755.

36. Kebrom TH, McKinley BA, Mullet JE. 2020. Shade signals alter the expression of circadian clock genes in newly-formed bioenergy sorghum internodes. Plant Direct 4: e00235.

37. Kim W-Y, Fujiwara S, Suh S-S, Kim J, Kim Y, Han L, David K, Putterill J, Nam HG, Somers DE. 2007. ZEITLUPE is a circadian photoreceptor stabilized by GIGANTEA in blue light. Nature 449: 356–360.

38. Kölling K, Thalmann M, Müller A, Jenny C, Zeeman SC. 2015. Carbon partitioning in Arabidopsis thaliana is a dynamic process controlled by the plants metabolic status and its circadian clock. Plant, Cell & Environment 38: 1965–1979.

39. Langfelder P, Horvath S. 2008. WGCNA: an R package for weighted correlation network analysis. BMC bioinformatics 9: 559.

40. Lembke CG, Nishiyama MY, Sato PM, de Andrade RF, Souza GM. 2012. Identification of sense and antisense transcripts regulated by drought in sugarcane. Plant Molecular Biology 79: 461–477.

41. Liu TL, Newton L, Liu M-J, Shiu S-H, Farré EM. 2016. A G-Box-Like Motif Is Necessary for Transcriptional Regulation by Circadian Pseudo-Response Regulators in Arabidopsis. Plant Physiology 170: 528–539.

42. MacKinnon KJ-M, Cole BJ, Yu C, Coomey JH, Hartwick NT, Remigereau M-S, Duffy T, Michael TP, Kay SA, Hazen SP. 2020. Changes in ambient temperature are the prevailing cue in determining Brachypodium distachyon diurnal gene regulation. New Phytologist 227: 1709– 1724.

43. Martínez-García JF, Huq E, Quail PH. 2000. Direct Targeting of Light Signals to a Promoter Element-Bound Transcription Factor. Science 288: 859–863.

44. Masclaux-Daubresse C, Valadier M-H, Carrayol E, Reisdorf-Cren M, Hirel B. 2002. Diurnal changes in the expression of glutamate dehydrogenase and nitrate reductase are involved in the C/N balance of tobacco source leaves. *Plant*, Cell & Environment 25: 1451–1462.

45. Matos DA, Cole BJ, Whitney IP, MacKinnon KJ-M, Kay SA, Hazen SP. 2014. Daily changes in temperature, not the circadian clock, regulate growth rate in Brachypodium distachyon. PloS One 9: e100072.

46. Matsuzaki J, Kawahara Y, Izawa T. 2015. Punctual transcriptional regulation by the rice circadian clock under fluctuating field conditions. The Plant Cell 27: 633–648.

47. McCormick AJ, Cramer MD, Watt DA. 2008. Regulation of photosynthesis by sugars in sugarcane leaves. Journal of Plant Physiology 165: 1817–1829.

48. Midmore DJ. 1980. Effects of photoperiod on flowering and fertility of sugarcane (Saccharum spp.). Field Crops Research 3: 65–81.

49. Mockler TC, Michael TP, Priest HD, Shen R, Sullivan CM, Givan SA, McEntee C, Kay SA, Chory J. 2007. The DIURNAL project: DIURNAL and circadian expression profiling, model-based pattern matching, and promoter analysis. Cold Spring Harbor Symposia on Quantitative Biology 72: 353–363.

50. Moore PH, Berding N. 2013. Flowering. In: Sugarcane: Physiology, Biochemistry, and Functional Biology. John Wiley & Sons, Ltd, 379–410.

51. Murphy RL, Klein RR, Morishige DT, Brady JA, Rooney WL, Miller FR, Dugas DV, Klein PE, Mullet JE. 2011. Coincident light and clock regulation of pseudoresponse regulator protein 37 (PRR37) controls photoperiodic flowering in sorghum. Proceedings of the National Academy of Sciences of the United States of America 108: 16469–16474.

52. Nagano AJ, Kawagoe T, Sugisaka J, Honjo MN, Iwayama K, Kudoh H. 2019. Annual transcriptome dynamics in natural environments reveals plant seasonal adaptation. Nature Plants 5: 74.

53. Nagano AJ, Sato Y, Mihara M, Antonio BA, Motoyama R, Itoh H, Nagamura Y, Izawa T. 2012. Deciphering and prediction of transcriptome dynamics under fluctuating field conditions. Cell 151: 1358–1369.

54. Nakamichi N, Kiba T, Henriques R, Mizuno T, Chua N-H, Sakakibara H. 2010. PSEUDO­RESPONSE REGULATORS 9, 7, and 5 are transcriptional repressors in the Arabidopsis circadian clock. The Plant Cell 22: 594–605.

55. Ni Z, Kim E-D, Ha M, Lackey E, Liu J, Zhang Y, Sun Q, Chen ZJ. 2009. Altered circadian rhythms regulate growth vigour in hybrids and allopolyploids. Nature 457: 327–331.

56. Nueda MJ, Tarazona S, Conesa A. 2014. Next maSigPro: updating maSigPro bioconductor package for RNA-seq time series. Bioinformatics 30: 2598–2602.

57. Panter PE, Muranaka T, Cuitun-Coronado D, Graham CA, Yochikawa A, Kudoh H, Dodd AN. 2019. Circadian Regulation of the Plant Transcriptome Under Natural Conditions. Frontiers in Genetics 10.

58. Preston JC, Kellogg EA. 2006. Reconstructing the Evolutionary History of Paralogous APETALA1/FRUITFULL-Like Genes in Grasses (Poaceae). Genetics 174: 421–437.

59. Pyl E-T, Piques M, Ivakov A, Schulze W, Ishihara H, Stitt M, Sulpice R. 2012. Metabolism and Growth in Arabidopsis Depend on the Daytime Temperature but Are Temperature-Compensated against Cool Nights. The Plant Cell 24: 2443–2469.

60. R Core Team. 2021. R: A language and environment for statistical computing.

61. Rao PS. 1977. Effects of Flowering on Yield and Quality of Sugarcane. Experimental Agriculture 13: 381–387.

62. Rawat R, Takahashi N, Hsu PY, Jones MA, Schwartz J, Salemi MR, Phinney BS, Harmer SL. 2011. REVEILLE8 and PSEUDO-REPONSE REGULATOR5 form a negative feedback loop within the Arabidopsis circadian clock. PLoS genetics 7: e1001350.

63. Reshef N, Fait A, Agam N. 2019. Grape berry position affects the diurnal dynamics of its metabolic profile. *Plant*, Cell & Environment 42: 1897–1912.

64. Roessner U, Luedemann A, Brust D, Fiehn O, Linke T, Willmitzer L, Fernie A. 2001. Metabolic profiling allows comprehensive phenotyping of genetically or environmentally modified plant systems. The Plant Cell 13: 11–29.

65. Shibaya T, Hori K, Ogiso-Tanaka E, Yamanouchi U, Shu K, Kitazawa N, Shomura A, Ando T, Ebana K, Wu J, et al. 2016. Hd18, Encoding Histone Acetylase Related to Arabidopsis FLOWERING LOCUS D, is Involved in the Control of Flowering Time in Rice. Plant & Cell Physiology 57: 1828–1838.

66. Song YH, Kubota A, Kwon MS, Covington MF, Lee N, Taagen ER, Laboy Cintrón D, Hwang DY, Akiyama R, Hodge SK, et al. 2018. Molecular basis of flowering under natural long-day conditions in Arabidopsis. Nature Plants 4: 824–835.

67. Steed G, Ramirez DC, Hannah MA, Webb AAR. 2021. Chronoculture, harnessing the circadian clock to improve crop yield and sustainability. Science 372.

68. Tanaka N, Itoh H, Sentoku N, Kojima M, Sakakibara H, Izawa T, Itoh J-I, Nagato Y. 2011. The COP1 Ortholog PPS Regulates the Juvenile–Adult and Vegetative–Reproductive Phase Changes in Rice. The Plant Cell 23: 2143–2154.

69. Webb AAR, Seki M, Satake A, Caldana C. 2019. Continuous dynamic adjustment of the plant circadian oscillator. Nature Communications 10: 550.

70. Weckwerth W, Wenzel K, Fiehn O. 2004. Process for the integrated extraction, identification and quantification of metabolites, proteins and RNA to reveal their co-regulation in biochemical networks. Proteomics 4: 78–83.

71. Yazdanbakhsh N, Sulpice R, Graf A, Stitt M, Fisahn J. 2011. Circadian control of root elongation and C partitioning in Arabidopsis thaliana. Plant, Cell & Environment 34: 877–894.

72. Zhao J, Huang X, Ouyang X, Chen W, Du A, Zhu L, Wang S, Deng XW, Li S. 2012. OsELF3-1, an Ortholog of Arabidopsis EARLY FLOWERING 3, Regulates Rice Circadian Rhythm and Photoperiodic Flowering. PLOS ONE 7: e43705.

